# BNP-Track: A framework for superresolved tracking

**DOI:** 10.1101/2023.04.03.535459

**Authors:** Ioannis Sgouralis, Lance W.Q. Xu, Ameya P. Jalihal, Nils G. Walter, Steve Pressé

## Abstract

Assessing dynamic processes at single molecule scales is key toward capturing life at the level of its molecular actors. Widefield superresolution methods, such as STORM, PALM, and PAINT, provide nanoscale localization accuracy, even when distances between fluorescently labeled single molecules (“emitters”) fall below light’s diffraction limit. However, as these superresolution methods rely on rare photophysical events to distinguish emitters from both each other and background, they are largely limited to static samples. In contrast, here we leverage spatiotemporal correlations of dynamic widefield imaging data to extend superresolution to simultaneous multiple emitter tracking without relying on photodynamics even as emitter distances from one another fall below the diffraction limit. We simultaneously determine emitter numbers and their tracks (localization and linking) with the same localization accuracy per frame as widefield superresolution does for immobilized emitters under similar imaging conditions (≈50 nm). We demonstrate our results for both *in cellulo* data and, for benchmarking purposes, on synthetic data. To this end, we avoid the existing tracking paradigm relying on completely or partially separating the tasks of emitter number determination, localization of each emitter, and linking emitter positions across frames. Instead, we develop a fully joint posterior distribution over the quantities of interest, including emitter tracks and their total, otherwise unknown, number within the Bayesian nonparametric paradigm. Our posterior quantifies the full uncertainty over emitter numbers and their associated tracks propagated from origins including shot noise and camera artefacts, pixelation, stochastic background, and out-of-focus motion. Finally, it remains accurate in more crowded regimes where alternative tracking tools cannot be applied.

## Introduction

Characterizing macromolecular assembly kinetics [16], quantifying intracellular biomolecular motility [71, 38, 39], or interrogating pairwise biomolecular interactions [88] requires accurate decoding of spatiotemporal processes at the single molecule level, *i*.*e*., high, nm spatial and rapid, often ms, temporal scales. These tasks ideally require superresolving positions of dynamic targets, typically fluorescently labeled molecules (light emitters), to tens of nanometer spatial resolution [51, 96, 5] and, when more than one target is involved, discriminating between signals from multiple targets simultaneously.

Assessments by means of fluorescence experiments at the required scales suffer from inherent limitations. Such limitations often arise from the diffraction limit of light, ≈250 nm in the visible range, below which conventional fluorescence techniques cannot achieve sufficient contrast between neighboring emitters of interest. To go beyond the limitations of conventional tools and achieve superresolution, contrast can be created through structured illumination [31], structured detection [41, 100, 99, 67], the photoresponse of fluorophore labels to excitation light [51, 96, 5, 79], or combinations thereof [34, 47, 101, 3].

Here we focus on widefield superresolution microscopy (SRM), which typically relies on fluorophore photodynamics to achieve superresolution. SRM is regularly used both *in vitro* [83, 42] and *in cellulo* [96, 5, 36, 95, 46, 65]. Specific widefield SRM image acquisition protocols, such as STORM [79], PALM [5], and PAINT [83], through their associated image analyses, decode positions of light emitters separated by distances below the diffraction limit, often down to the resolution of tens of nanometers [79, 5]. These widefield SRM protocols can be broken down into three conceptual steps: (i) specimen preparation; (ii) imaging; and (iii) computational processing of the acquired images (frames). The success of Step (iii) is ensured by both Steps (i)-(ii). In particular, in Step (i) engineered fluorophores are selected that enable the desired photodynamics; *e*.*g*., photoswitching in STORM [79], photo-activation/bleaching in PALM [5], or fluorophore binding/unbinding in PAINT [83]. Step (ii) is then performed over extended periods, while awaiting rare photophysical (or binding-unbinding) events to manifest and for sufficient photons to be collected to achieve superresolved localizations in Step (iii). For well-isolated bright spots, Step (iii) achieves superresolved localization [44, 51, 96] while accounting for effects such as light diffraction, resulting in spot sizes of roughly twice 0.61*λ/*NA (the Rayleigh diffraction limit), set by the emitter wavelength (*λ*), the numerical aperture (NA) of the microscope’s objective [74], the camera and its photon shot noise and spot pixelization.

In this work, we show that computation can be used to overcome reliance both on photophysics in Step (i) and on a long acquisition time in Step (ii), which not only limits widefield SRM largely to spatiotemporally fixed samples, but can also induce sample photodamage. In fact, while a moving emitter’s distribution, or smearing, of its photon budget over multiple frames and pixels is a net disadvantage in the implementation of Step (iii) above, we demonstrate– conversely–that such a distribution of the photon budget in both space and time provides information that can be leveraged to superresolve emitter tracks, determine emitter numbers, and help discriminate targets from their neighbors, even in the complete absence of photophysical processes (fig. 1).

**Figure 1:**
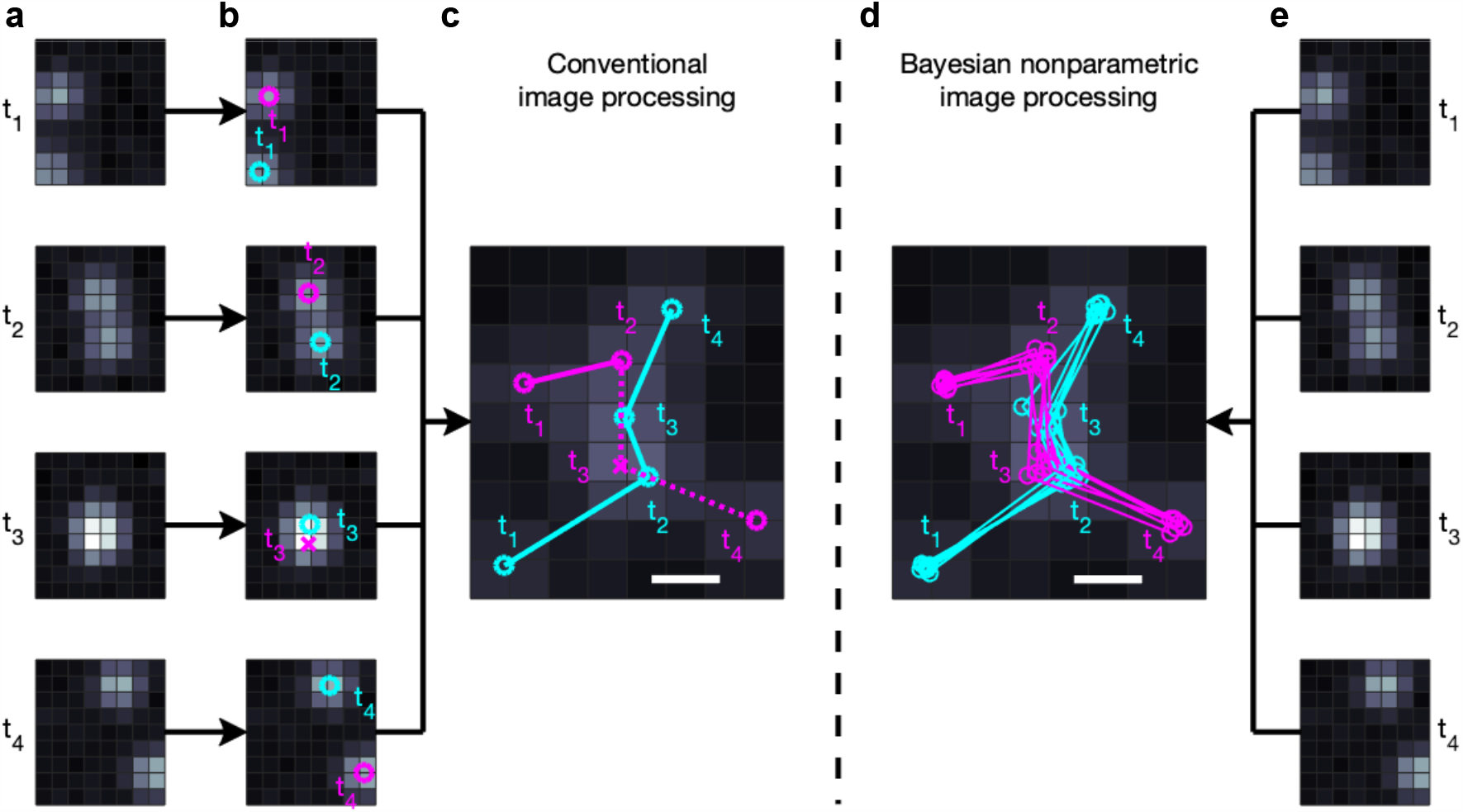
Conceptual comparison between widely available tracking frameworks and BNP-Track. **a** and **e**, Four frames from a dataset showing two emitters. **b**, Existing tracking approaches [91, 40, 105, 57, 11, 103, 18, 30, 52, 106, 37, 9] either completely or partially separate the task of first identifying and then localizing light emitters in the field of view of each frame independently. c, Conventional approaches then link emitter positions across frames. **d**, Our nonparametric approach (BNP-Track) simultaneously determines the number of emitters, localizes them and links their positions across frames. In **b-d**, circles denote correctly identified emitters, and crosses denote missed emitters. In **c** and **d**, scale bars indicate a distance equal to the nominal diffraction limit given by the Rayleigh diffraction limit 0.61λ/NA.

Although captured in more detail in the framework put forward in Methods and Supporting Information, here, we briefly highlight how our tracking framework, Bayesian nonparametrics (BNP)-Track, differs fundamentally from conventional tracking tools that determine emitter numbers, localize emitters, and link emitter locations as sequential steps. In the language of Bayesian statistics, resolving emitter tracks as well as emitter numbers amounts to constructing the probability distribution, ℙ (links, locations, emitter numbers|data), which reads as “the joint posterior probability distribution of emitter numbers, locations, and links given a dataset”. The best set of emitter number and tracks are those globally maximizing this probability distribution. Without further approximation, Bayes’ theorem allows us to decompose this probability distribution as this product:

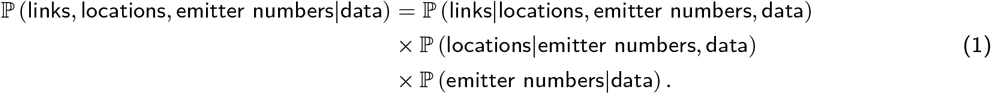

Single particle tracking (SPT) tools that perform emitter number determination, emitter localization, and linking in separate steps [93, 6, 81, 7, 72, 27, 24, 89, 40, 85, 12, 18, 17, 63, 103, 102, 30, 76, 52, 54, 19, 2, 84, 53, 90, 8, 37, 77, 87, 50, 49, 9, 29, 104, 13, 15, 1, 66, 22, 61, 56, 58, 11, 68, 60, 78, 25, 14, 4, 59, 55, 91, 86], many reviewed here [14, 96, 80, 10, 51], invariably approximate the maximization of the joint distribution as a serial maximization of three terms. This process often involves additional approximations, such as using ℙ (links|locations) to approximate ℙ (links|locations, emitter numbers, data). Approximations such as these are acceptable for well-isolated and in-focus emitters, though they have fundamentally limited our ability to superresolve emitters especially as they move closer than light’s diffraction limit. In contrast, BNP-Track avoids all such approximations and leverages all sources of information through the joint posterior to draw samples as well as maximize the posterior probability distribution, yielding superresolved emitter tracks.

The overall input to BNP-Track includes both raw image sequences and known information on the imaging system, including the microscope optics and camera electronics as further detailed in Methods. Using Bayesian nonparametrics, we provide a means of estimating unknowns including the number of emitters and their associated tracks. We demonstrate BNP-Track on experimental SPT data, detailing how the simultaneous determination of emitter numbers and tracks can be computationally achieved. We also benchmark BNP-Track’s performance against a well-established diffraction-limited tracking tool, TrackMate [91], to which we must confer some advantage as direct comparison is not possible since existing tools do not simultaneously learn emitter numbers and associated tracks. More detailed comparisons are presented in Results.

## Results

To demonstrate our approach, we use BNP-Track to first analyze single mRNA molecules diffusing in live U-2 OS cells imaged under single-plane HILO illumination [92] on a fluorescence microscope as previously described [71, 69, 20], but using a beamsplitter to divide the single-color signal onto two cameras (fig. 2). The dual-camera setup allows us to test for consistency of BNP-Track’s emitter number and track determination across cameras. In subsequent tests, we employ noise-overlaid synthetic data for which the ground truth is known [91, 14] (figs. 3 to 5), and finally challenge BNP-Track with experimental data of crowded emitters (fig. 6).

**Figure 2:**
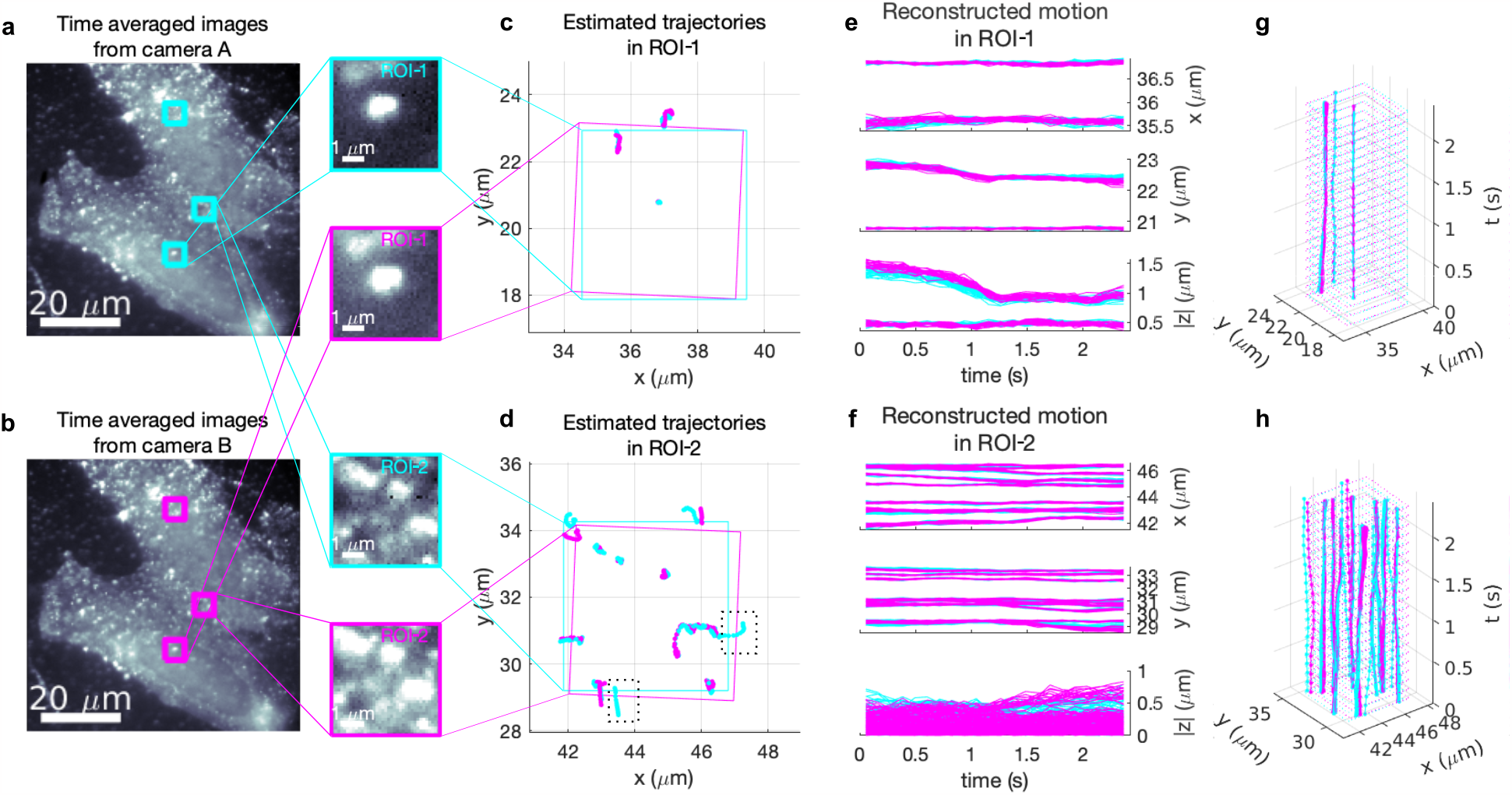
Testing BNP-Track’s performances on two 5 μm-wide regions of interest with different emitter densities based on an experimental dataset from fluorophore-labeled mRNA molecules diffusing in live U-2 OS cells onto a dual-camera microscope. **a** and **b**, For convenience only, we show time-averages of all 22 frames analyzed from camera A and B, respectively. The selected ROIs are boxed. We zoom in on ROI-1 and ROI-2 in the subpanels. The remaining region, ROI-3, is only highlighted and analyzed later in the text (fig. 6). **c** and **d**, Estimated tracks within the selected ROIs from both cameras with solid boxes indicating the corresponding ROIs after image registration. **e** and **f**, Reconstructed time courses for individual tracks from the selected ROIs. The dotted boxes in d highlight two emitter tracks only detected by camera A. **g**, Time course reconstruction by combining (**e**, top) and (**e**, middle). **h**, Time course reconstruction by combining (**f**, top) and (**f**, middle).

**Figure 3:**
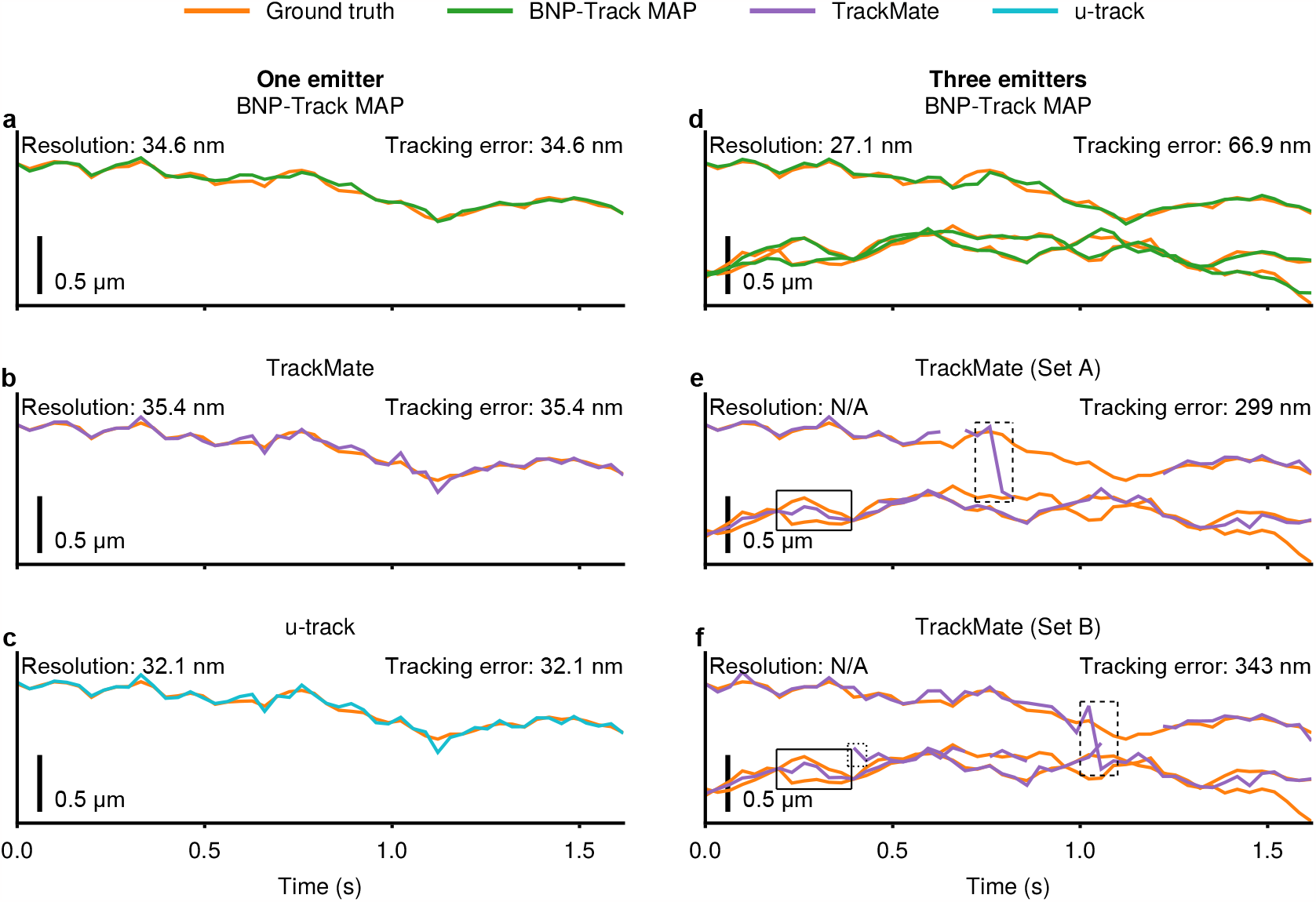
A comparison of tracking performance among BNP-Track, TrackMate, and u-track using two synthetic datasets with one emitter (Supplementary Video 6) and three emitters (Supplementary Video 7) in the y coordinate. See fig. A.3 for the same figure but with the x coordinate. **a**, BNP-Track’s MAP estimate compared with the one-emitter ground truth. **b**, TrackMate’s estimate compared with the one-emitter ground truth. Set A and Set B are equivalent in this case. **c**, u-track’s estimate compared with the one-emitter ground truth. d, BNP-Track’s MAP estimates compared with the three-emitter ground truth. **e** and **f**, TrackMate’s estimates compared with the three-emitter ground truth, respectively. In (**d-f**), the top ground truth track is the same as the ground truth in (**a-c**). The boxed regions in (**e**) and (**f**) highlight where TrackMate performs relatively poorly.

**Figure 4:**
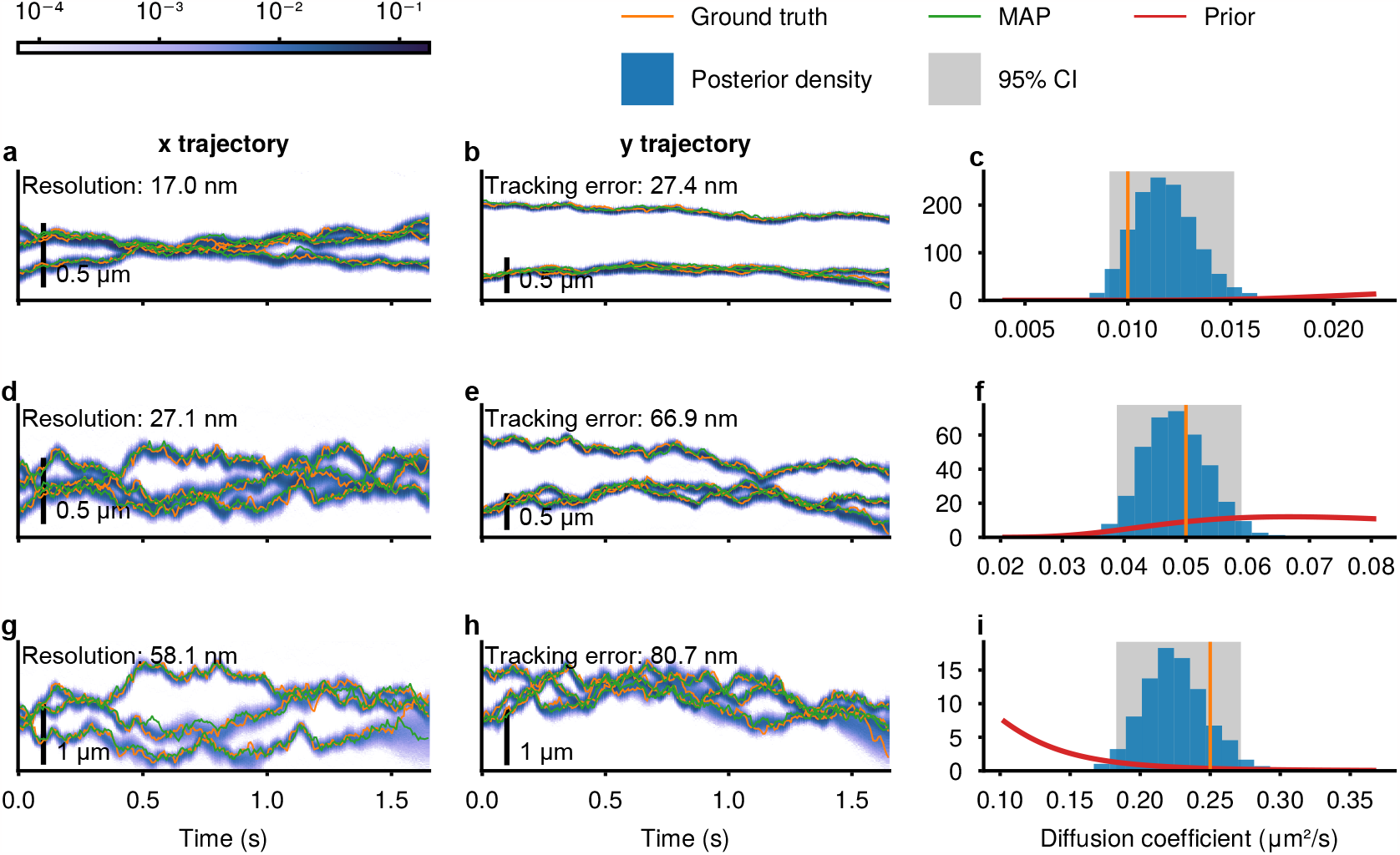
BNP-Track’s performance under various diffusion coefficients. **a-c**, BNP-Track’s analysis for Supplementary Video 8 with 0.01 μm^2^s^−1^. **d-f**, BNP-Track’s analysis for Supplementary Video 7 with 0.05 μm^2^s^−1^. **g-i**, BNP-Track’s analysis for are from Supplementary Video 9 with 0.25 μm^2^s^−1^. Credible intervals (CIs) are depicted as shaded regions (which we refer to as CI bands), with darker shading indicating a higher level of confidence.

**Figure 5:**
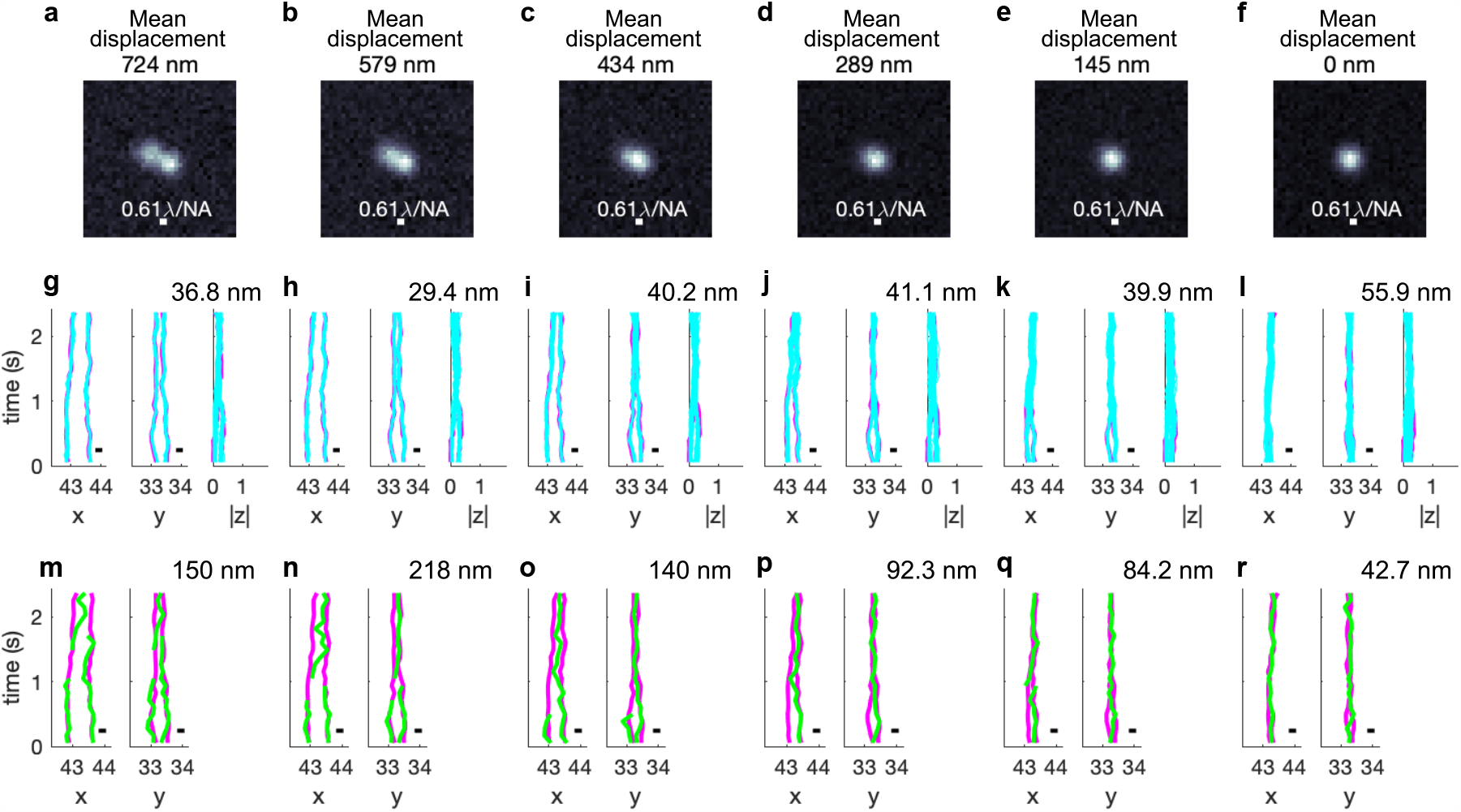
Benchmarks of BNP-Track regarding the mean displacement between simulated tracks. TrackMate tracks (Set B) are also provided for comparison. **a-f**, The time-averaged images for the synthetic scenarios (Supplementary Videos 11-16). The mean displacements are 724 nm (**a**), 579 nm (**b**), 434 nm (**c**), 289 nm (**d**), 145 nm (**e**), and 0 nm (**f**). **g-l**, BNP-Track’s reconstructed tracks for all coordinates. Reconstructed tracks are in cyan and the ground truths are in magenta. Lateral tracking errors are listed at the top right of each panel. **m-r**, TrackMate’s estimated tracks (green) of the same datasets with the ground truths (magenta). Lateral tracking errors are listed at the top right of each panel, with missing segments excluded. Also, no axial results are plotted in (**m-r**) as TrackMate does not provide axial tracks for 2D images.

As emitters move in 3D, it is possible, indeed helpful in more accurate lateral localization, for BNP-Track to estimate emitter axial distance (|*z*|) from the in-focus plane from 2D images using the width of the emitter’s point spread function (PSF) sensitivity to axial distance [107]. For this reason, while axial distance from the in-focus plane is always less accurately determined than lateral positions, we nonetheless report BNP-Track’s axial estimates for experimental data in figs. 2 and 6.

**Figure 6:**
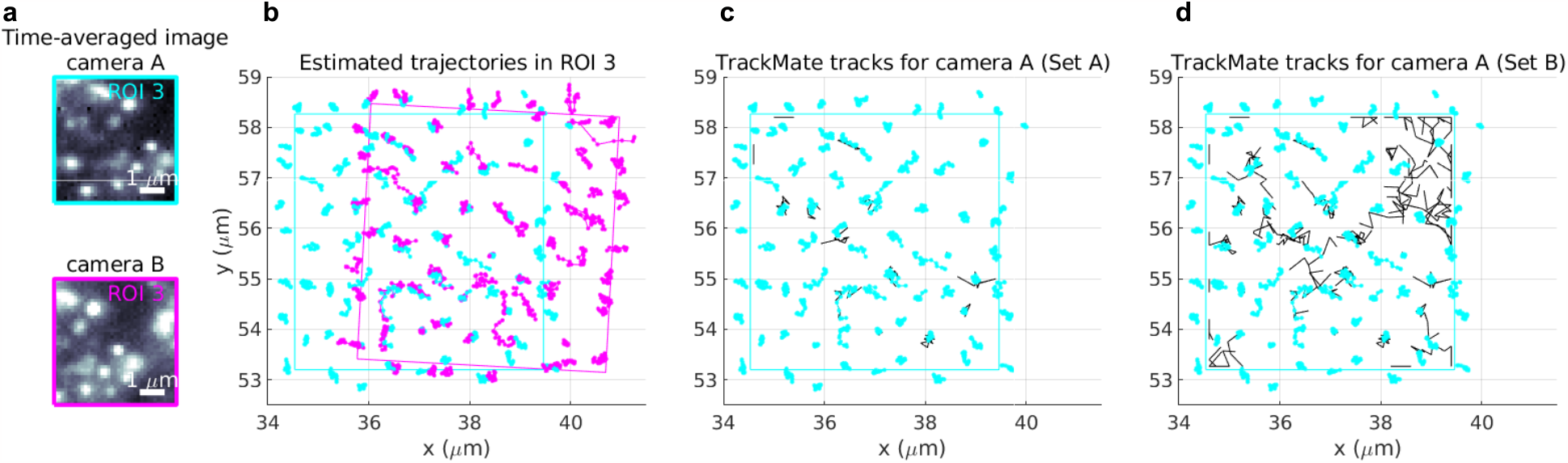
BNP-Track’s performance in increasingly crowded environments. **a**, Time-averaged images over 22 frames from both cameras show that there are many diffraction-limited emitters within the ROI analyzed here, ROI-3. **b**, BNP-Track’s tracking results after registering fields of view. **c**, A comparison between BNP-Track’s result and TrackMate’s result with a high localization quality threshold (Set A) for the data from camera A. TrackMate tracks are in black; see [91] for the definition of quality. **d**, A comparison between BNP-Track’s result and TrackMate’s result with a low localization quality threshold (Set B) for the data from camera A. TrackMate tracks are in black.

Prior to showing results of BNP-Track, we make note of an important feature of Bayesian inference. Developed within the Bayesian paradigm [94, 28, 97], BNP-Track not only provides point estimates over unknown quantities of interest, such as numbers of emitters and their associated tracks, but also distributions over them, which includes uncertainty. As we cannot easily visualize the output of multidimensional posteriors over all candidate emitter numbers and associated tracks, we often report estimates for emitters which coincide with the number of emitters that maximize the posterior termed *maximum a posteriori* (MAP) point estimates. Then, having determined the MAP number of emitters, we collect their associated tracks in figures such as figs. 2 and 6.

### BNP-Track superresolves sparse single-emitter tracks in cellulo

To assess the success of BNP-Track, as no direct ground truth is available for tracks from experimental SPT data, we employ two separate cameras behind a beamsplitter. Using image registration (to correct for camera misalignment), we independently process two datasets for subsequent comparison and error estimation knowing that, in principle, both cameras should have the same tracks (our ground truth) even though the noise realizations on both cameras is different and particles may move closer to each other than the nominal diffraction limit.

The agreement between tracks across both cameras within their field of view can be quantified according to a pairing distance metric [14]. Briefly, the distance between two tracks is defined using the gated Euclidean distance given by

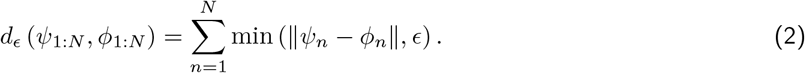

Here, *ϵ* is called the gate value, *N* is the total number of frames, and ∥*ψ*_*n*_ − *ϕ*_*n*_∥ denotes the Euclidean distance between the emitter positions *θ*_*n*_ and *ϕ*_*n*_ at time *t*_*n*_ in both cameras. If a track fails to localize any emitter in a particular frame, the distance at that frame is considered to be *ϵ*. The pairing distance between two sets of tracks is defined as the minimum total gated Euclidean distance among all possible track pairings between the sets. When comparing BNP-Track’s tracking results for both cameras in the dual-camera setup, we use an infinite value for *ϵ* as, by design, BNP-Track only outputs tracks with no missing segments. Additionally, we introduce the concept of tracking error, calculated as the pairing distance divided by the product of the number of frames and the number of emitters.

In figs. 2a and 2b, we show, for illustrative purposes alone, time averages of a sequence of 22 successive frames spanning ≈2.5 s of real time in both detection channels. In data processing, we analyze the underlying frames without averaging. All raw data are provided in Supporting Information (Supplementary Videos 1-4).

From these frames, we track well separated or dilute emitters, *i*.*e*., whose PSFs always are well separated in space, in a 5 *μ*m wide square region of interest (ROI), named ROI-1. Figure 2a also zooms in on ROI-1. The BNP-Track derived track estimates are shown in fig. 2c, while in fig. 2e all samples from the posterior are superposed. Due to fundamental optical limitations, we cannot determine whether the emitter’s axial position lies above or below the in-focus plane, so we only report the absolute value of the emitters’ axial position. As evident from fig. 2c, BNP-Track successfully identifies and localizes the same tracks within the selected ROI in the two parallel camera datasets of ROI-1 despite different noise realizations on each camera and background. Of note, the two square ROI-1’s are rotated relative to one another based on image registration in post-processing. fig. 2e shows tracks of all well separated emitters identified within this field of view. We note that despite the fact that the center of the PSF of the emitter near the top of ROI-1 in fig. 2c is outside ROI-1 for both cameras in all 22 frames, it is still surprisingly independently picked up in the analysis of the data from both cameras. However, we exclude this unique track outside the field of view from fig. 2e and further analysis.

Using the metric defined above for fig. 2e, we report a tracking error (pairing distance averaged over the number of frames) of 73 nm in the lateral direction. Consequently, BNP-Track’s average error from the underlying ground truth is one half of the tracking error, or about 37 nm, consistent with prior superresolution values for immobilized targets [51, 96, 5].

As BNP-Track provides estimates of the lateral as well as the magnitude of the axial emitter position (see details in Methods), we can also assess the full tracking error in 3D. This results in a 3D pairing distance of 97 nm and thus a ≈48 nm tracking error from ground truth. Having shown that we can track emitters in a dilute regime similar to ROI-1 of figs. 2c and 2d, we next analyze a more challenging ROI, ROI-2, (figs. 2e and 2f) where emitter PSFs now occasionally overlap.

As before, for illustrative purposes alone, in figs. 2a and 2b we show time averages of a sequence of 22 successive frames spanning ≈2.5 s of real time. Figures 2d, 2f and 2h reflect the same information described for ROI-1 except that they are for ROI-2. Just as in ROI-1, BNP-Track can track emitters even when they diffuse away from a camera’s field of view. Using the pairing distance to quantify the tracking error in the lateral direction between the two cameras, for ROI-2 the tracking error is slightly higher at 128 nm with an error from the ground truth of about 64 nm, which remains under the nominal diffraction limit of 231 nm. Additionally, the tracking error in 3D is now slightly elevated compared to the one from ROI-1, to 159 nm, resulting in an error from ground truth of about 80 nm.

BNP-Track also estimates other dynamical quantities, including the background photon fluxes (photon per unit area per unit time), the emitter brightness (photon per unit time), the diffusion coefficient, and the number of emitters. Estimates for these quantities are summarized in fig. A.1. It is worth noting that figs. A.1b and A.1c show that both the system’s background flux and emitter brightness vary substantially over time, making ROI-2 more challenging. Despite the agreement between tracks deduced from both cameras below light’s diffraction limit, discrepancies in some quantities (such as diffusion coefficient in fig. A.1d) highlight the sensitivity of these quantities to small track differences below light’s diffraction limit. Similarly, small discrepancies in the emitter brightness estimates (fig. A.1c) may be induced by minute dissimilarities in the optical path leading to each camera.

Finally, the number of emitters detected in the two cameras are different (fig. A.1e), with the additional tracks detected by camera A highlighted by dotted boxes in fig. 2d. This is unsurprising for three reasons. First, the two cameras have slightly different fields of view. Second, as highlighted by the dotted boxes in fig. 2d, a significant portion of the two extra tracks lie outside the fields of view of either cameras and thus are difficult tracks to detect under any circumstance. Third, as background noise can be mathematically modeled by out-of-focus emitters, and since two cameras draw slightly different conclusions on background photon emission rates and emitter brightnesses (figs. A.1b and A.1c), this may also naturally lead to slightly different estimates of the number of emitters especially out-of-focus or out of the field of view. That is, BNP-Track not only detects in-focus emitters, but uses what it learns from in-focus emitters to extrapolate outside the field of view or the in-focus plane. In such regions the number of photons used by BNP-Track to draw inferences on tracks is naturally limited.

### Benchmarking BNP-Track with synthetic data

We next validate our method by employing synthetically generated data where ground truth tracks are known. To ensure realistic data, we adopt the procedure outlined in Methods and Supporting Information for data generation. The parameters (NA, pixel size, frame rate, diffusion coefficient, emitter brightness, and background photon flux) utilized in generating synthetic data are either identical or similar to the experiments discussed earlier. All parameter values are specified in table B.1. By using simulated data and the accompanying ground truth tracks, we evaluate the performance of BNP-Track in two ways. First, we compare BNP-Track’s tracking accuracy to an established SPT tool built upon contest leading tracking methods [14], TrackMate [91]. Second, we test BNP-Track’s robustness across different parameter regimes.

However, comparing BNP-Track to other SPT methods in a fair and direct manner poses significant challenges. One challenge is due to the absence of any other existing method that can simultaneously estimate emitter number alongside the associated tracks (and with diffusion coefficient, time-dependent emitter brightness, and time-dependent background photon flux). Consequently, throughout the entire manuscript, we give a major advantage to existing tools that we compare to BNP-Track by: (i) manually tuning the parameters of these tools to have them best match the ground truth emitter numbers and locations; and (ii) asking BNP-Track to estimate parameters (diffusion coefficient and background photon flux) from the data whilst the ground truth values for these same parameters are used to tune competing methods to optimize their performance. See Methods for exactly how the aforementioned parameters are provided to existing methods. As we will show, even providing competing methods significant advantages, BNP-Track still exceeds the resolution of existing tools and yields reduced error rates (percentage of wrong links).

In comparing to existing tools, an important point arises from the fact that there exists no single metric to assess whether tracks obtained using conventional SPT tools are good or not. In the absence of a quantitative numerical criterion such as a posterior value, we rely on pre-selected metrics, *e*.*g*., tracks with minimal spurious detections or tracks with the fewest missed links (termed false negatives). Also, although it is generally preferable to have tracks with no false positives (spurious detections or tracks) and no false negatives (missed detections or tracks), it is often challenging for competing methods to achieve both simultaneously. This difficulty arises because most methods lack the capability to set frame-specific thresholds. Consequently, when false negatives and false positives cannot be reduced to zero simultaneously for a conventional SPT tool, we use SPT tools to output two sets of tracks from data: Set A prioritizes minimizing false positives during the localization (spot detection) phase and subsequently minimizes false negatives during the linking phase. In contrast, Set B focuses on minimizing false negatives during the localization phase and then addresses false positives during the linking phase.

To fairly compare SPT tools to BNP-Track, we continue using the pairing distance and tracking error metrics previously employed. In addition, we use a finite gate value (*ϵ*, see eq. (2)) of five pixels (approximately 665 nm). This gate value allows us to benchmark SPT tools that otherwise face challenges in localizing emitters within specific frames (*i*.*e*., have missing segments) using a defined threshold. Nonetheless, the pairing distance metric has a limitation that must be addressed: it does not directly reflect localization accuracy since it penalizes linking errors. To better capture localization accuracy, we introduce another metric called localization resolution, or simply resolution, that does not consider linking error. Instead of pairing tracks, localization resolution independently pairs emitter positions in each frame without employing a gate value. Therefore, localization resolution is undefined if the tracks compared do not have the same number of emitters in any given frame. Also, if there are no linking errors, false positives, or false negatives, tracking error and localization resolution are equivalent.

### Comparison with TrackMate

We start with the simplest possible case of only one emitter in the field of view. In this case, both BNP-Track (fig. 3a) and TrackMate (fig. 3b) successfully track the single emitter throughout the entire video with similar resolution (34.6 nm vs. 35.4 nm). In order to further demonstrate that conventional SPT tools would typically work under this simple scenario, we also include the track obtained with u-track [40] using the point source particle detection process (fig. 3c), which again achieves a similar resolution at 32.1 nm. We note that in figs. 3a to 3c, resolution is equal to tracking error as there are no missing segments or incorrect links in tracks.

Having established that both BNP-Track and TrackMate work well for a simple case, we move on to a more complicated dataset with three emitters. Predictably, as we begin encountering PSF overlap in multiple emitters for the three-emitter dataset (see fig. A.2a and Supplementary Video 7), BNP-Track and TrackMate’s performances diverge (we note that u-track’s performance is not shown here as the current FOV is too small for running its algorithm). While BNP-Track remains capable of tracking all three emitters throughout the entire video with a resolution of 27.1 nm from the MAP tracks, even as these fall below the diffraction limit in frames 2 to 13 and 34 to 47 (see fig. A.2a), several issues arise for the TrackMate tracks (figs. 3e and 3f). For instance, diffraction-limited emitters get interpreted as one emitter (solid boxes), incorrect links with large jumps (dashed boxes), and spurious detections (the dotted box) from TrackMate Set B. These issues indicate that TrackMate can no longer resolve the emitters in this dataset, and hence no resolution is calculated. As for the tracking error between the ground truth tracks and each SPT method’s output: BNP-Track’s MAP tracks have a tracking error of 66.9 nm while both TrackMate track sets yield tracking errors no less than 300 nm.

Three additional points should be noted here: (i) As illustrated in fig. 3d, BNP-Track’s tracking error is larger than its resolution, which is due to the linking error. Nevertheless, we posit that this is not a major concern since it only occurs near frame 35 where two emitters are laterally less than 40 nm apart (see fig. A.2a) while one of them is out-of-focus (300 nm away from the in-focus plane). Under this circumstance, these two emitters are very difficult to distinguish. (ii) Even though the three-emitter dataset contains the same track as the one-emitter dataset, BNP-Track actually achieves a better resolution in the more complicated three-emitter dataset. We attribute this finding to the fact that BNP-Track leverages all spatiotemporal correlation as detailed in the next section. (iii) TrackMate Set B does a little better than Set A in estimating emitter numbers though increasing the overall tracking error.

More quantitative comparisons generated by the Tacking Performance Measures plugin [14] in Icy [21] are provided in tables B.2 and B.3 in Supporting Information.

In this section, we test BNP-Track’s robustness regarding two quantities of interest, diffusion coefficient and the spatial separation between emitters.

BNP-Track remains robust in the analysis of image sequences generated with different diffusion coefficients, as shown in fig. 4. Here, BNP-Track accurately tracks all emitters, as determined by localization resolution, and determines the correct diffusion coefficient, even when the diffusion coefficient changes by a factor of 25 from the slowest to the fastest case. As observed in fig. 4, the distribution of emitter positions becomes broader as the diffusion coefficient increases. This is supported by the 95% confidence intervals of the localization resolutions, which range from (15.2 to 18.8) nm in the first row, to (22.8 to 27.3) nm in the second row, to (47.8 to 65.4) nm in the third row. One major factor contributing to this trend is motion blur introduced by increasing diffusion coefficients. Another important factor is that faster diffusing species have less time to remain within the field of view or move away from the in-focus plane, resulting in fewer informative frames (frames with sufficient detected photons).

As a general framework, when needed, BNP-Track can also be extended to incorporate faster-diffusing species. Briefly, we do so by estimating multiple positions for each emitter within each camera exposure period. See appendix G.4 for a mathematical explanation and, in fig. A.5, we test this extension using a synthetic dataset, Supplementary Video 10, consisting of an emitter diffusing at 1 *μ*m^2^s^−1^ with camera exposure being 30 ms.

Next, in fig. 5, we test how closely two emitters can come together while retaining BNP-Track’s ability to enumerate the number of emitters and track them. To this end, a pair of estimated tracks from ROI-2 of fig. 2 were used as ground truth for the simulation of synthetic data (shown in figs. 5a to 5f) using the same parameters. Then, the mean displacement between emitters is gradually decreased. Figures 5g to 5l show reconstructed tracks and comparison with the ground truth. Remarkably, the tracking error estimated from the total pairing distance remains ≈40 nm in 2D and slightly increases (by ≈50 nm) in 3D throughout the synthetic scenarios, below the diffraction limit.

As seen in figs. 3e and 3f, existing tracking methods, such as TrackMate, can fail to resolve emitters with separations close to or below the diffraction limit. To further demonstrate this point, we show the results from TrackMate Set B (which better estimates emitter numbers). When the mean displacement between emitters remains well above the lateral diffraction limit (figs. 5a to 5c), TrackMate Set B can be tuned to resolve two emitters (figs. 5m to 5o) albeit with missing track segments. These missing track segments are attributed to out-of-focus emitters. This is most clearly evidenced by noting that in figs. 5a to 5c one emitter’s average intensity distribution is broader. More quantitatively, this observation is because this emitter, as seen in figs. 5g to 5i, has a |*z*| position at a distance 200 nm from the in-focus plane. Once the mean displacement is close to or below the lateral diffraction limit (figs. 5d to 5f), even Set B can no longer be tuned to resolve two emitters in any frame. In contrast, by leveraging spatiotemporal information, BNP-Track successfully tracks out-of-focus emitters. Additional comparisons (using quantitative performance metrics) are again provided in tables B.4 to B.9.

### BNP-Track’s performance in increasingly crowded environments

So far, we have evaluated the performance of BNP-Track on two distinct ROIs from an experimental dataset and confirmed its resolution using synthetic data. To further test the limits of BNP-Track, we analyzed a densely-packed ROI, ROI-3 (Supplementary Videos 17 and 18), selected from the same data set; see fig. 2a.

Here in fig. 6a, similarly to figs. 2a and 2b, we first present time-averaged images from both cameras. These images reveal that ROI-3 contains tens of closely positioned as well as out-of-focus emitters. Furthermore, within ROI-3, cameras A and B observe slightly different fields of view, offset by approximately 1.5 *μ*m and rotated by 5^*°*^. To provide an assessment of BNP-Track’s performance to comparison with that of TrackMate, we selected the in-focus emitters in the overlapping region and calculated the pairing distance between tracks. We define “in-focus” emitters as emitters whose estimated *z* position is within 150 nm of the in-focus (*z* = 0) plane. The results show a tracking error derived from this pairing distance of approximately 136.4 nm, which corresponds to a tracking error of 68.2 nm compared to the ground truth. These results are consistent with the performance for ROI-2 in fig. 2.

As illustrated in figs. 6a and 6b, ROI-3 presents a challenge for emitter number estimation due to crowding and overlapping PSFs, as well as larger number of out-of-focus emitters. These features pose serious challenges to conventional tracking tools relying on manually setting thresholds to fix the number of emitters especially dim or out-of-focus ones. To demonstrate this point, we again use TrackMate to analyze the dataset from ROI-3. The analysis results for camera A are illustrated in figs. 6c and 6d. Specifically, for fig. 6c we set a relative high localization quality (Set A) threshold at 5 to minimize spurious detection, resulting in a total of 18 tracks.

Each of these tracks can be paired with one of BNP-Track’s 78 emitter tracks using the Tracking Performance Measures tool [14] in Icy [21], with the maximum pairing distance set at two pixels or 266 nm (which is close to the nominal diffraction limit of 231 nm). Even with a high localization threshold, TrackMate produces significantly fewer emitter tracks than BNP-Track due to out-of-focus dim emitters and difficulties arising from overlapping

PSFs. In contrast, for fig. 6d, we lower the quality threshold in TrackMate to 0 to detect more emitters (Set B), which increases the total number of TrackMate tracks to 64. Using the same pairing distance threshold, 41 of these tracks can be paired with a subset of BNP-Track’s emitter tracks. However, there are also 23 spurious tracks contaminating further analysis. Similar results are also obtained for data from camera B, see figs. A.6a and A.6b. Furthermore, in figs. A.6c and A.6d we show that, given the same image registration as used in fig. 6b, TrackMate does not yield matching tracks for both cameras. These results underscore the importance of inferring the number of emitters simultaneously while tracking rather than pre-calibrating emitter numbers.

## Discussion

We present here an image processing framework, BNP-Track, that superresolves particles *in cellulo* without the need for complex fluorophore photodynamics. Our framework analyzes continuous image measurements from diffraction-limited light emitters throughout image acquisition. BNP-Track thereby extends the scope of widefield SRM by leveraging spatiotemporal information encoded across frames and pixels. Additionally, BNP-Track unifies many existing approaches to localization microscopy and single-particle tracking and extends beyond them by simultaneously and self-consistently estimating emitter numbers.

By operating in three interlaced stages (preparation, imaging, processing) existing approaches to widefield SRM estimate locations of individual static emitters with a generally accepted resolution of ≈50 nm or less [51, 96, 5]. Such resolution for widefield applications is a significant improvement relative to the diffraction limit of conventional microscopy, ≈250 nm. Although our processing framework cannot lift the limitations imposed by optics, nor eliminate the degradation induced by noise, we show that our framework can significantly extend our ability to estimate emitter numbers and tracks from existing images with uncertainty both for easier in-focus cases where emitters are well separated to more challenging cases where emitters are crowded, move out-of-focus as well as cases where emitters appear partly out of the field of view. In particular, because our framework provides full distributions over unknowns, it readily computes error bars (often termed credible intervals in a Bayesian setting) associated with emitter numbers warranted by images (*e*.*g*., figs. A.1 and A.4), localization events for both isolated emitters and emitters closer than light’s diffraction limit, and other parameters including diffusion coefficients.

It is clear that many tracking scenarios provide challenges for any tracking tool by limiting the information encoded in the data available for analysis. These scenarios include out-of-focus motion, crowded environments, or even detector saturation. Although quantifying precisely when BNP-Track fails depends on the specifics of any given circumstance, BNP-Track leverages broad spatiotemporal information typically eliminated by separating the tracking task in modular steps by traditional tools as highlighted in Introduction. BNP leverages all information available by modeling the entire process from emitter motion to detector output simultaneously and self-consistently, thus maximizing the amount of information extracted from individual frames. As such, when BNP-Track fails to track for a particular system setting, conventional tracking methods typically fail earlier, for example as shown in fig. 5, indicating the need for an alternative experimental protocol.

The analysis of a field of view like those shown in Results (5 *μ*m *×* 5 *μ*m or about 1500 pixels, and 22 frames) requires about 300 min of computational time on an average laptop. The computational cost of our analysis scales linearly with the number of frames, total pixel number, and total number of emitters. Larger scale applications are within the realm of existing computational capacity; though additional algorithmic improvements and computational optimization may help speed up execution time. Despite its higher computational cost compared to current single-particle tracking methods, we argue two points. Firstly, as demonstrated in figs. 3, 5 and 6, cheaper conventional tracking methods not only fail to surpass the diffraction limit, but they also do not learn emitter numbers. Yet learning emitter numbers is especially critical in correctly linking emitter locations across frames, especially in crowded environments. Secondly, our method’s time of execution is primarily computational wall-time as BNP-Track is unsupervised and largely free from manual tuning. This stands in contrast to methods such as TrackMate used in the generation of figs. 3, 5 and 6 requiring manual tuning and thresholding for proper execution and, even so, remain diffraction limited.

While we have focused on the conceptual basis for how to beat the diffraction limit computationally for moving targets, with more extensive changes to our processing framework the methods herein can also be adapted to accommodate specialized illumination modalities including TIRF [23] and light-sheet [75], or even multi-color imaging. All of these adaptations are compatible with Bayesian nonparametric tracking, which relies on the joint assessment of three steps characteristic of traditional single-particle tracking (emitter number determination, emitter localization, and track linking). Along these same lines, in Methods, we made many common modeling choices and used parameters typical for experiments. For example, we used an EMCCD camera model and assumed a Gaussian PSF. Other choices can be made by simply changing the mathematical form of the camera model or the PSF, provided these assume known, pre-calibrated forms. None of these changes break the conceptual framework of BNP-Track.

Similarly, another assumption made in this study is that BNP-Track considers only a Brownian motion model, and one may question its performance when emitters evolve according to alternative motion models. However, throughout Results, we have demonstrated that BNP-Track yields accurate tracking results consistent across two cameras for an experimental dataset with an unknown emitter motion model, despite assuming Brownian motion. This may suggest that BNP-Track remains robust under other motion models. We may thus conceive of the Brownian motion model instead as simply providing a motivation to use Gaussian transition probabilities, following from the central limit theorem, between locations across frames. What is more, if we have reason to believe that a specific motion model is warranted that may not be accommodated by Gaussian transition probabilities, we may also incorporate this change into our framework.

Currently, post-processing tools are frequently utilized to extract useful information from single-particle tracks, such as diffusion coefficients and diffusive states. These tools range from simple approaches, such as mean square displacement, to more complex methods, such as Spot-On [32] and SMAUG [43]. Since our framework produces tracks, these tracks can be analyzed by these tools and our ability to produce full distributions over tracks may also be helpful in estimating errors over post-processed parameters. It is also conceivable that our output could be used as a training set for neural networks [48] or be used to make predictions of molecular tracks in dense environments [26], such as fig. 6, previously considered outside the scope of existing tools.

Taken together, the framework we present is the first proof-of-principle demonstration that computation feasibly achieves superresolution of evolving targets by avoiding the modular approximations of the existing tracking paradigm, which thus far has limited tracking to dilute and in-focus samples.

## Methods

### Image processing

As we demonstrate in Results, our analysis goal is the determination of the probability distribution termed the posterior, 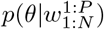. In this distribution, we use *θ* to gather the unknown quantities of interest, for instance emitter tracks and photon emission rates, and 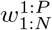 to gather the data under processing, for instance timelapse images. Below we present how this distribution is derived as well as its underlying assumptions.

To facilitate the presentation, we first present a detailed formulation of the physical processes involved in the formation of the acquired images necessary in quantitative analysis discussed earlier. This formulation captures microscope optics and camera electronics and can be further modified to accommodate more specialized imaging setups. In the formulation, the unknowns of interest are encoded by parameters. Next, we present the mathematical tools needed to estimate values for such unknown parameters. That is, we address the core challenge in SRM arising from the unknown number of emitters and their associated tracks. To overcome the challenge of estimating emitter numbers, we apply Bayesian nonparametrics. Our approach is different from likelihood-based approaches, currently employed in localization microscopy, and is ultimately what allows us to relax SRM’s photodynamical requirements.

As several of the notions involved in our description are stochastic, for example parameters with unknown values and random emitter dynamics, we use probabilistic descriptions. Although our notation is standard to the statistics community, we provide an introduction more appropriate for a broader audience in Supporting Information.

### Model description

Our starting point consists of image measurements obtained in a SRM experiment denoted by 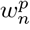, where subscripts *n* = 1, …, *N* indicate the exposures and superscripts *p* = 1, …, *P* indicate pixels. For example, 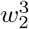 denotes the raw image value, typically reported in ADU or counts and stored in TIFF format, measured in pixel 3 during the second exposure. Similarly, 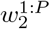 denotes every image value (*i*.*e*., entire frame) measured during the second exposure. Since the image values are related to the specimen under imaging, our goal from now on is to develop a mathematical model encoding the physical processes relating system imaged with the acquired measurements.

### Noise

Recorded images mix electronic signals that depend only stochastically on an average amount of incident photons [73, 45, 35, 33]. For commercially available cameras, the overall relationship, from incident photons to recorded images, is linear and contaminated with multiplicative noise that results from shot noise, amplification and readout. Our formulation below applies to image data acquired with cameras of EMCCD type, as commonly used in superresolution imaging [51, 35], though the expression below can be modified to accommodate other detector architectures. Here, in our formulation

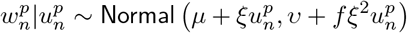

where 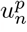 is the average number of photons incident on pixel *p* during exposure *n*. The parameter *f* is a camera dependent excess noise factor and *ξ* is the overall gain that combines the effects of quantum efficiency, preamplification, amplification and quantization. The values of *μ, υ, ξ*, and *f* are specific to the camera acquiring the images of interest and their values can be calibrated as we describe in Supporting Information.

### Pixelization

As shot-noise is already captured, 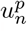 depends deterministically on the underlying photon flux

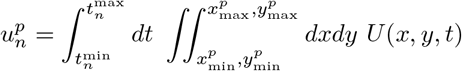

where 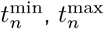 mark the integration time of the *n*^th^ exposure; 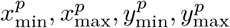 mark the region monitored by pixel *p*; and *U* (*x, y, t*) is the photon flux at position *x, y* and time *t*. We detail our spatiotemporal frames of reference in Supporting Information.

### Optics

We model *U* (*x, y, t*) as consisting of background *U*_back_(*x, y, t*) and fluorophore photon contributions, *i*.*e*., flux, from every imaged light emitter 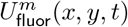. These are additive

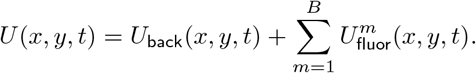

Specifically, for the latter, we consider a total of *B* emitters that we label with *m* = 1, …, *B*. Each of our emitters is characterized by a position *X*^*m*^(*t*), *Y* ^*m*^(*t*), *Z*^*m*^(*t*), all of which may change through time. Here, we use uppercase letters *X, Y*, and *Z* for random variables and lowercase letters *x, y*, and *z* for general variables (realizations of the corresponding random variables). Since the total number *B* of imaged emitters is a critical unknown quantity, in the next section we describe how we modify the flux *U* (*x, y, t*) to allow for a variable number of emitters and in Supporting Information we describe how this flux is related to *X*^*m*^(*t*), *Y* ^*m*^(*t*), *Z*^*m*^(*t*).

### Model inference

The quantities we wish to estimate, for example the positions *X*^1:*B*^(*t*), *Y* ^1:*B*^(*t*), *Z*^1:*B*^(*t*), are unknown variables in the preceding formulation. The total number of such variables depends upon the number of imaged emitters *B*, which in SRM remains unknown thereby prohibiting processing of the images under the flux *U* (*x, y, t*). Since *B* has such subtle effect, we modify our formulation to make it compatible with the nonparametric paradigm of Data Analysis that allows for processing under an unspecified number of variables [98, 64, 94, 28, 62].

In particular, following the nonparametric latent feature paradigm [94, 28], we introduce indicator parameters *b*^*m*^ that adopt only values 0 or 1 and recast *U* (*x, y, t*) in the form

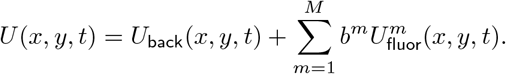

Specifically, with the introduction of indicators, we increase the number of emitters represented in our model from *B* to a number *M > B* that may be arbitrarily large. The critical advantage is that now the total number of model emitters *M* may be set before processing in contrast with the total number of actual emitters *B* that remains unknown. With this formulation we infer the values of *b*^1:*M*^ during processing, simultaneously with the other parameters of interest. This way, we can actively recruit, *i*.*e*., *b*^*m*^ = 1, or discard, *i*.*e*., *b*^*m*^ = 0, light emitters consistently avoiding under/overfitting. Following image processing, our analysis recovers the total number of imaged emitters by the sum 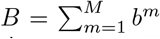 and the positions of the emitters *X*^*m*^(*t*), *Y* ^*m*^(*t*), *Z*^*m*^(*t*) by the estimated positions of those model emitters with *b*^*m*^ = 1. However, a side effect of introducing *M* is that our analysis results may depend on the particular value chosen. To relax this dependence, we use a specialized nonparametric prior on *b*^*m*^ that we describe in detail in S upporting I nformation . This prior specifically allows for image processing at the formal limit *M → ∞*.

Our overall formulation also includes additional parameters, for example background photon fluxes and fluorophore brightness, that may or may not be of immediate interest. To provide a flexible computational scheme, that works around both unknown types, *i*.*e*., parametric and nonparametric, and also allows for future extensions, we adopt a Bayesian approach in which we prescribe prior probability distributions on every unknown parameter beyond just the indicators *b*^*m*^. These priors, combined with the preceding formulation, lead to the posterior probability distribution 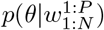, where *θ* gathers every unknown, on which our results rely. We describe the full posterior distribution and its evaluation in S upporting I nformation .

### TrackMate, u-track, and Tracking Performance Measure

Besides its widespread use, ongoing maintenance and updates, and being built upon leading methods in Ref. [14], we opt for TrackMate [91] specifically because it combines various localization and linking methods, as well as multiple thresholding options.

To generate tracks for comparison in R esults, we first export simulated videos as TIFF files and import them in Fiji [82] v1.54b for analysis with TrackMate v7.9.2. As part of implementing TrackMate, the Laplacian of Gaussian detector with sub-pixel localization and the linear assignment problem (LAP) mathematical framework [40] are used in spot detection. Spots are then filtered based on quality, contrast, sum intensity, and radius (set based on the actual PSF size used in data simulation). For the LAP tracker, we allow gap closing and tune, based on diffusion coefficients, the parameters for max (inter-frame) distance, max frame gap, and number of spots in tracks to find the best tracks. No extra feature penalties are added. All aforementioned parameters are tuned with the aim of minimizing tracking error.

u-track [40] is employed for the sake of additional comparison, following the same tuning process described earlier. As an advantage to u-track, we provide u-track with the background emission (termed “absolute background” in their manual), which we instead learn from the simulated data using BNP-Track.

The benchmarks in tables B.2 and B.3 were created using the Tracking Performance Measure plugin in Icy [21] 2.4.3.0. To generate these benchmarks, we exported the TrackMate track, the BNP-Track MAP estimates, and ground truth tracks as XML files. All tracks were imported into Icy’s TrackManager using the “Import TrackMate track file” feature, and the Tracking Performance Measure plugin was initiated using the “add Track Processor” option. The only required input for this plugin is the “maximum distance between detections”, which we kept at a default value of five pixels.

### Image acquisition Experimental timelapse images

Fluorescence timelapse images of U-2 OS cells injected with chemically labeled firefly luciferase mRNAs were acquired simultaneously on two cameras. Cell culture and handling of U-2 OS cells prior to injections were performed as previously described [70]. Firefly luciferase mRNAs were *in vitro* transcribed, capped and polyadenylated, and a variable number of Cy3 dyes were non specifically added to the polyA tail using CLICK chemistry [20]. Cells were injected with a solution of Cy3 labeled mRNAs and Cascade Blue labeled 10 kDa dextran (Invitrogen D1976) using a Femtojet pump and Injectman NI2 micromanipulator (Eppendorf) at 20 hPa for 0.1 s with 20 hPa compensation pressure. Successfully injected cells were identified by the presence of a fluorescent dextran and were imaged 30 minutes post injection. Cells were continuously illuminated with the 532 nm laser in HILO mode, and Cy3 fluorescence was collected using a 60x 1.49 NA oil objective. Images were captured simultaneously on two Andor X-10 EMCCD cameras by using a 50:50 beamsplitter, with a 100 ms exposure time.

### Synthetic timelapse images

We acquire the validation and benchmarking data through standard computer simulations. We start from ground truth as specified in the captions of figs. 3 to 5, A.4 and A.5 and then added noise with values we estimated from the experimental timelapse images according to appendix F.

## Acknowledgements

We acknowledge NIH NIGMS (R01GM130745) for supporting early efforts in nonparametrics and tracking, NIH NIGMS (R01GM134426) for supporting single photon efforts, NIH MIRA (R35GM148237) entitled “Toward high spatiotemporal resolution models of single molecules for in cellulo applications. We also acknowledge R01GM122803 and R35GM131922 (to N.G.W.) for enabling experimental data acquisition, as well as NSF MRI-ID grant DBI-0959823 (to N.G.W.) for seeding the Single Molecule Analysis in Real-Time (SMART) Center, whose Single Particle Tracker TIRFM equipment was used for acquiring experimental tracking data with support from J.D. Hoff. A.P.J. was partially supported by NIH T-32-GM007315.

## References

[1] A. Agrawal, M. Gupta, A. Veeraraghavan, and S. G. Narasimhan. Optimal coded sampling for temporal super-resolution. In 2010 IEEE Computer Society Conference on Computer Vision and Pattern Recognition, page 599, 2010.

[2] C. M. Anderson, G. N. Georgiou, I. Morrison, G. Stevenson, and R. J. Cherry. Tracking of cell surface receptors by fluorescence digital imaging microscopy using a charge-coupled device camera. low-density lipoprotein and influenza virus receptor mobility at 4 degrees c. J. Cell. Sci., 101(2):415, 1992.

[3] F. Balzarotti, Y. Eilers, K. C. Gwosch, A. H. Gynnå, V. Westphal, F. D. Stefani, J. Elf, and S. W. Hell. Nanometer resolution imaging and tracking of fluorescent molecules with minimal photon fluxes. Science, 355(6325):606, 2017.

[4] A. O. Barden, A. S. Goler, S. C. Humphreys, S. Tabatabaei, M. Lochner, M.-D. Ruepp, T. Jack, J. Simonin, A. J. Thompson, J. P. Jones, and J. A. Brozik. Tracking individual membrane proteins and their biochemistry: The power of direct observation. Neuropharmacology, 98:22, 2015.

[5] E. Betzig, G. H. Patterson, R. Sougrat, O. W. Lindwasser, S. Olenych, J. S. Bonifacino, M. W. Davidson, J. Lippincott-Schwartz, and H. F. Hess. Imaging intracellular fluorescent proteins at nanometer resolution. Science, 313(5793):1642, 2006.

[6] S. Bonneau, M. Dahan, and L. Cohen. Single quantum dot tracking based on perceptual grouping using minimal paths in a spatiotemporal volume. IEEE Trans. Image Process., 14(9):1384, 2005.

[7] B. C. Carter, G. T. Shubeita, and S. P. Gross. Tracking single particles: A user-friendly quantitative evaluation. Phys. Biol., 2(1):60, Mar. 2005.

[8] I. Casuso, J. Khao, M. Chami, P. Paul-Gilloteaux, M. Husain, J.-P. Duneau, H. Stahlberg, J. N. Sturgis, and S. Scheuring. Characterization of the motion of membrane proteins using high-speed atomic force microscopy. Nat. Nanotechnol., 7(8):525, 2012.

[9] K. Celler, G. P. van Wezel, and J. Willemse. Single particle tracking of dynamically localizing tata complexes in streptomyces coelicolor. Biochem. Biophys. Res. Commun., 438(1):38, 2013.

[10] H.-J. Cheng, C.-H. Hsu, C.-L. Hung, and C.-Y. Lin. A review for cell and particle tracking on microscopy images using algorithms and deep learning technologies. Biomed. J., 45(3):465, 2022.

[11] N. Chenouard, I. Bloch, and J.-C. Olivo-Marin. Multiple hypothesis tracking for cluttered biological image sequences. IEEE Trans. Pattern Anal. Mach. Intell., 35(11):2736, 2013.

[12] N. Chenouard, s. Bloch, and J.-C. Olivo-Marin. Feature-aided particle tracking. In 2008 15th IEEE International Conference on Image Processing, page 1796, 2008.

[13] N. Chenouard, A. Dufour, and J.-C. Olivo-Marin. Tracking algorithms chase down pathogens. Biotechnol. J., 4(6):838–845, 2009.

[14] N. Chenouard, I. Smal, F. De Chaumont, M. Maška, I. F. Sbalzarini, Y. Gong, J. Cardinale, C. Carthel, S. Coraluppi, M. Winter, et al. Objective comparison of particle tracking methods. Nat. Methods, 11(3):281, 2014.

[15] M. Chertkov, L. Kroc, F. Krzakala, M. Vergassola, and L. Zdeborová. Inference in particle tracking experiments by passing messages between images. Proc. Natl. Acad. Sci. U.S.A., 107(17):7663, 2010.

[16] I. I. Cisse, I. Izeddin, S. Z. Causse, L. Boudarene, A. Senecal, L. Muresan, C. Dugast-Darzacq, B. Hajj, M. Dahan, and X. Darzacq. Real-time dynamics of rna polymerase ii clustering in live human cells. Science, 341(6146):664, 2013.

[17] S. Coraluppi and C. Carthel. Recursive track fusion for multi-sensor surveillance. Inf. Fusion, 5(1):23, 2004.

[18] S. Coraluppi and C. Carthel. Multi-stage multiple-hypothesis tracking. J. Adv. Inf. Fusion, 6(1):57, 2011.

[19] J. C. Crocker and D. G. Grier. Methods of digital video microscopy for colloidal studies. J. Colloid Interface Sci., 179(1):298–310, 1996.

[20] T. C. Custer and N. G. Walter. In vitro labeling strategies for in cellulo fluorescence microscopy of single ribonucleoprotein machines. Protein Sci., 26(7):1363, 2017.

[21] F. De Chaumont, S. Dallongeville, N. Chenouard, N. Hervé, S. Pop, T. Provoost, V. Meas-Yedid, P. Pankajakshan, T. Lecomte, Y. Le Montagner, et al. Icy: an open bioimage informatics platform for extended reproducible research. Nat. Methods, 9(7):690, 2012.

[22] A. Dufour, R. Thibeaux, E. Labruyere, N. Guillen, and J.-C. Olivo-Marin. 3-D active meshes: Fast discrete deformable models for cell tracking in 3-D time-lapse microscopy. IEEE Trans. Image Process., 20(7):1925, 2011.

[23] K. N. Fish. Total internal reflection fluorescence (tirf) microscopy. Curr. Protoc. Cytom., 50(1):12, 2009.

[24] E. Fox, E. Sudderth, and A. Willsky. Hierarchical Dirichlet processes for tracking maneuvering targets. In 2007 10th International Conference on Information Fusion, 2007.

[25] E. B. Fox, M. C. Hughes, E. B. Sudderth, and M. I. Jordan. Joint modeling of multiple time series via the beta process with application to motion capture segmentation. Ann. Appl. Stat., 8(3):1281, 2014.

[26] N. Galvanetto, M. T. Ivanović, A. Chowdhury, A. Sottini, M. Nüesch, D. Nettels, R. Best, and B. Schuler. Ultrafast molecular dynamics observed within a dense protein condensate. bioRxiv, pages 2022–12, 2022.

[27] A. Genovesio, T. Liedl, V. Emiliani, W. Parak, M. Coppey-Moisan, and J. Olivo-Marin. Multiple particle tracking in 3-D+t microscopy: Method and application to the tracking of endocytosed quantum dots. IEEE Trans. Image Process., 15:1062, 2006.

[28] Z. Ghahramani. Probabilistic machine learning and artificial intelligence. Nature, 521(7553):452, 2015.

[29] W. Godinez, M. Lampe, S. Wörz, B. Müller, R. Eils, and K. Rohr. Deterministic and probabilistic approaches for tracking virus particles in time-lapse fluorescence microscopy image sequences. Med. Image Anal., 13(2):325, 2009.

[30] W. J. Godinez, M. Lampe, R. Eils, B. Müller, and K. Rohr. Tracking multiple particles in fluorescence microscopy images via probabilistic data association. In 2011 IEEE International Symposium on Biomedical Imaging: From Nano to Macro, page 1925. IEEE, 2011.

[31] M. G. Gustafsson. Surpassing the lateral resolution limit by a factor of two using structured illumination microscopy. J. Microsc., 198(2):82, 2000.

[32] A. S. Hansen, M. Woringer, J. B. Grimm, L. D. Lavis, R. Tjian, and X. Darzacq. Robust model-based analysis of single-particle tracking experiments with spot-on. Elife, 7:e33125, 2018.

[33] K. B. Harpsøe, M. I. Andersen, and P. Kjægaard. Bayesian photon counting with electron-multiplying charge coupled devices (EMCCDs). Astron. Astrophys., 537:A50, 2012.

[34] S. W. Hell and J. Wichmann. Breaking the diffraction resolution limit by stimulated emission: stimulated-emission-depletion fluorescence microscopy. Opt. Lett., 19(11):780, 1994.

[35] M. Hirsch, R. J. Wareham, M. L. Martin-Fernandez, M. P. Hobson, and D. J. Rolfe. A stochastic model for electron multiplication charge-coupled devices – from theory to practice. PLoS One, 8(1):1, 2013.

[36] S. J. Holden, T. Pengo, K. L. Meibom, C. Fernandez Fernandez, J. Collier, and S. Manley. High throughput 3d super-resolution microscopy reveals caulobacter crescentus in vivo z-ring organization. Proc. Natl. Acad. Sci. U.S.A., 111(12):4566, 2014.

[37] M. Husain, T. Boudier, P. Paul-Gilloteaux, I. Casuso, and S. Scheuring. Software for drift compensation, particle tracking and particle analysis of high-speed atomic force microscopy image series. J. Mol. Recognit., 25(5):292, 2012.

[38] A. P. Jalihal, S. Pitchiaya, L. Xiao, P. Bawa, X. Jiang, K. Bedi, A. Parolia, M. Cieslik, M. Ljungman, A. M. Chinnaiyan, et al. Multivalent proteins rapidly and reversibly phase-separate upon osmotic cell volume change. Mol. Cell, 79(6):978–990, 2020.

[39] A. P. Jalihal, A. Schmidt, G. Gao, S. R. Little, S. Pitchiaya, and N. G. Walter. Hyperosmotic phase separation: Condensates beyond inclusions, granules and organelles. J. Biol. Chem., 296, 2021.

[40] K. Jaqaman, D. Loerke, M. Mettlen, H. Kuwata, S. Grinstein, S. L. Schmid, and G. Danuser. Robust single-particle tracking in live-cell time-lapse sequences. Nat. Methods, 5(8):695, 2008.

[41] S. Jazani, L. W. Xu, I. Sgouralis, D. P. Shepherd, and S. Pressé. Computational proposal for tracking multiple molecules in a multifocus confocal setup. ACS photonics, 9(7):2489, 2022.

[42] R. Jungmann, M. S. Avendaño, J. B. Woehrstein, M. Dai, W. M. Shih, and P. Yin. Multiplexed 3d cellular super-resolution imaging with dna-paint and exchange-paint. Nat. Methods, 11(3):313, 2014.

[43] J. D. Karslake, E. D. Donarski, S. A. Shelby, L. M. Demey, V. J. DiRita, S. L. Veatch, and J. S. Biteen. Smaug: Analyzing single-molecule tracks with nonparametric bayesian statistics. Methods, 193:16, 2021.

[44] I. M. Khater, I. R. Nabi, and G. Hamarneh. A review of super-resolution single-molecule localization microscopy cluster analysis and quantification methods. Patterns, 1(3):100038, 2020.

[45] Z. Kilic, I. Sgouralis, W. Heo, K. Ishii, T. Tahara, and S. Pressé. Extraction of rapid kinetics from smfret measurements using integrative detectors. Cell Rep. Phys. Sci., 2(5):100409, 2021.

[46] J. Kim, J. Y. Kim, S. Jeon, J. W. Baik, S. H. Cho, and C. Kim. Super-resolution localization photoacoustic microscopy using intrinsic red blood cells as contrast absorbers. Light Sci. Appl., 8(1):103, 2019.

[47] T. A. Klar, S. Jakobs, M. Dyba, A. Egner, and S. W. Hell. Fluorescence microscopy with diffraction resolution barrier broken by stimulated emission. Proc. Natl. Acad. Sci. U.S.A., 97(15):8206, 2000.

[48] P. Kowalek, H. Loch-Olszewska, and J. Szwabiński. Classification of diffusion modes in single-particle tracking data: Feature-based versus deep-learning approach. Phys. Rev. E, 100(3):032410, 2019.

[49] T.-C. Ku, Y.-N. Huang, C.-C. Huang, D.-M. Yang, L.-S. Kao, T.-Y. Chiu, C.-F. Hsieh, P.-Y. Wu, Y.-S. Tsai, and C.-C. Lin. An automated tracking system to measure the dynamic properties of vesicles in living cells. Microsc. Res. Tech., 70(2):119, 2007.

[50] T.-C. Ku, L.-S. Kao, C.-C. Lin, and Y.-S. Tsai. Morphological filter improve the efficiency of automated tracking of secretory vesicles with various dynamic properties. Microsc. Res. Tech., 72(9):639, 2009.

[51] A. Lee, K. Tsekouras, C. Calderon, C. Bustamante, and S. Pressé. Unraveling the thousand word picture: An introduction to super-resolution data analysis. Chem. Rev., 117(11):7276, 2017.

[52] L. Liang, H. Shen, P. De Camilli, and J. S. Duncan. Tracking clathrin coated pits with a multiple hypothesis based method. In Medical Image Computing and Computer-Assisted Intervention–MICCAI 2010: 13th International Conference, Beijing, China, September 20-24, 2010, Proceedings, Part II 13, page 315. Springer, 2010.

[53] D. G. Lowe. Distinctive image features from scale-invariant keypoints. Int. J. Comput. Vis., 60:91, 2004.

[54] K. E. Magnusson and J. Jaldén. A batch algorithm using iterative application of the viterbi algorithm to track cells and construct cell lineages. In 2012 9th IEEE International Symposium on Biomedical Imaging (ISBI), page 382. IEEE, 2012.

[55] V. Maroulas and A. Nebenführ. Tracking rapid intracellular movements: A Bayesian random set approach. Ann. Appl. Stat., 9(2):926, 2015.

[56] E. Meijering, O. Dzyubachyk, and I. Smal. Chapter nine - methods for cell and particle tracking. In P. M. conn, editor, Imaging and Spectroscopic Analysis of Living Cells, volume 504 of Methods in Enzymology, page 183. Academic Press, 2012.

[57] E. Meijering, O. Dzyubachyk, and I. Smal. Methods for cell and particle tracking. Meth. Enzymol., 504:183, 2012.

[58] X. Michalet and A. Berglund. Optimal diffusion coefficient estimation in single-particle tracking. Phys. Rev. E, 85:061916, 2012.

[59] N. Monnier, Z. Barry, H. Y. Park, K.-C. Su, Z. Katz, B. P. English, A. Dey, K. Pan, I. M. Cheeseman, R. H. Singer, and M. Bathe. Inferring transient particle transport dynamics in live cells. Nat. Methods, 12(9):838, 2015.

[60] A. D. Mont, C. P. Calderon, and A. B. Poore. A new computational method for ambiguity assessment of solutions to assignment problems arising in target tracking. In O. E. Drummond, editor, Signal and Data Processing of Small Targets 2014, volume 9092, page 159. International Society for Optics and Photonics, SPIE, 2014.

[61] U. Mudenagudi, S. Banerjee, and P. K. Kalra. Space-time super-resolution using graph-cut optimization. IEEE Trans. Pattern Anal. Mach. Intell., 33(5):995, 2011.

[62] P. Müller, F. A. Quintana, A. Jara, and T. Hanson. Bayesian nonparametric data analysis, volume 1. Springer, 2015.

[63] J.-C. Olivo-Marin. Extraction of spots in biological images using multiscale products. Pattern Recognit., 35(9):1989–1996, 2002.

[64] P. Orbanz and Y. W. Teh. Encyclopedia of machine learning, volume 1, chapter Bayesian Nonparametric Models. Springer Science & Business Media, 2010.

[65] J. Otterstrom, A. Castells-Garcia, C. Vicario, P. A. Gomez-Garcia, M. P. Cosma, and M. Lakadamyali. Super-resolution microscopy reveals how histone tail acetylation affects dna compaction within nucleosomes in vivo. Nucleic Acids Res., 47(16):8470, 2019.

[66] H. Y. Park, A. R. Buxbaum, and R. H. Singer. Chapter 18 - single mrna tracking in live cells. In N. G. Walter, editor, Single Molecule Tools: Fluorescence Based Approaches, Part A, volume 472 of Methods in Enzymology, page 387. Academic Press, 2010.

[67] E. P. Perillo, Y.-L. Liu, K. Huynh, C. Liu, C.-K. Chou, M.-C. Hung, H.-C. Yeh, and A. K. Dunn. Deep and high-resolution three-dimensional tracking of single particles using nonlinear and multiplexed illumination. Nat. Commun., 6(1):7874, 2015.

[68] F. Persson, M. Lindén, C. Unoson, and J. Elf. Extracting intracellular diffusive states and transition rates from single-molecule tracking data. Nat. Methods, 10(3):265, 2013.

[69] S. Pitchiaya, L. A. Heinicke, T. C. Custer, and N. G. Walter. Single molecule fluorescence approaches shed light on intracellular rnas. Chem. Rev., 114(6):3224, 2014.

[70] S. Pitchiaya, V. Krishnan, T. C. Custer, and N. G. Walter. Dissecting non-coding rna mechanisms in cellulo by single-molecule high-resolution localization and counting. Methods, 63(2):188, 2013.

[71] S. Pitchiaya, M. D. Mourao, A. P. Jalihal, L. Xiao, X. Jiang, A. M. Chinnaiyan, S. Schnell, and N. G. Walter. Dynamic recruitment of single rnas to processing bodies depends on rna functionality. Mol. Cell, 74(3):521, 2019.

[72] V. Racine, A. Hertzog, J. Jouanneau, J. Salamero, C. Kervrann, and J. Sibarita. Multiple-target tracking of 3D fluorescent objects based on simulated annealing. In 3rd IEEE International Symposium on Biomedical Imaging: Nano to Macro, 2006., page 1020, 2006.

[73] S. Ram, P. Prabhat, J. Chao, E. Sally Ward, and R. J. Ober. High accuracy 3d quantum dot tracking with multifocal plane microscopy for the study of fast intracellular dynamics in live cells. Biophys. J., 95(12):6025, 2008.

[74] L. Rayleigh. XXXI. investigations in optics, with special reference to the spectroscope. Lond. Edinb. Dubl. Phil. Mag., 8(49):261, 1879.

[75] E. G. Reynaud, J. Peychl, J. Huisken, and P. Tomancak. Guide to light-sheet microscopy for adventurous biologists. Nat. Methods, 12(1):30, 2015.

[76] J. Rink, E. Ghigo, Y. Kalaidzidis, and M. Zerial. Rab conversion as a mechanism of progression from early to late endosomes. Cell, 122(5):735, 2005.

[77] P. J. Rousseeuw and A. M. Leroy. Robust regression and outlier detection. John wiley & sons, 2005.

[78] D. J. Rowland and J. S. Biteen. Top-hat and asymmetric gaussian-based fitting functions for quantifying directional single-molecule motion. ChemPhysChem, 15(4):712, 2014.

[79] M. J. Rust, M. Bates, and X. Zhuang. Sub-diffraction-limit imaging by stochastic optical reconstruction microscopy (STORM). Nat. Methods, 3(10):793, 2006.

[80] M. J. Saxton and K. Jacobson. Single-particle tracking:applications to membrane dynamics. Annu. Rev. Biophys., 26(1):373, 1997. PMID: 9241424.

[81] I. Sbalzarini and P. Koumoutsakos. Feature point tracking and trajectory analysis for video imaging in cell biology. J. Struct. Biol., 151(2):182, 2005.

[82] J. Schindelin, I. Arganda-Carreras, E. Frise, V. Kaynig, M. Longair, T. Pietzsch, S. Preibisch, C. Rueden, S. Saalfeld, B. Schmid, J.-Y. Tinevez, D. J. White, V. Hartenstein, K. Eliceiri, P. Tomancak, and A. Cardona. Fiji: an open-source platform for biological-image analysis. Nat. Methods, 9(7):676, 2012.

[83] J. Schnitzbauer, M. T. Strauss, T. Schlichthaerle, F. Schueder, and R. Jungmann. Super-resolution microscopy with DNA-PAINT. Nature protocols, 12(6):1198, 2017.

[84] G. Seisenberger, M. U. Ried, T. Endress, H. Buning, M. Hallek, and C. Brauchle. Real-time single-molecule imaging of the infection pathway of an adeno-associated virus. Science, 294(5548):1929, 2001.

[85] A. Sergé, N. Bertaux, H. Rigneault, and D. Marguet. Dynamic multiple-target tracing to probe spatiotemporal cartography of cell membranes. Nat. Methods, 5(8):687, Aug. 2008.

[86] I. Sgouralis, A. Nebenführ, and V. Maroulas. A bayesian topological framework for the identification and reconstruction of subcellular motion. SIAM J. Imaging Sci., 10(2):871, 2017.

[87] K. Shafique and M. Shah. A noniterative greedy algorithm for multiframe point correspondence. IEEE Trans. Pattern Anal. Mach. Intell., 27(1):51, 2005.

[88] M. Shayegan, R. Tahvildari, K. Metera, L. Kisley, S. W. Michnick, and S. R. Leslie. Probing inhomogeneous diffusion in the microenvironments of phase-separated polymers under confinement. J. Am. Chem. Soc., 141(19):7751, 2019.

[89] I. Smal, K. Draegestein, N. Galjart, W. Niessen, and E. Meijering. Particle filtering for multiple object tracking in dynamic fluorescence microscopy images: Application to microtubule growth analysis. IEEE Trans. Med. Imaging, 27(6):789, 2008.

[90] R. E. Thompson, D. R. Larson, and W. W. Webb. Precise nanometer localization analysis for individual fluorescent probes. Biophys. J., 82(5):2775, 2002.

[91] J.-Y. Tinevez, N. Perry, J. Schindelin, G. M. Hoopes, G. D. Reynolds, E. Laplantine, S. Y. Bednarek, S. L. Shorte, and K. W. Eliceiri. Trackmate: An open and extensible platform for single-particle tracking. Methods, 115:80–90, 2017. Image Processing for Biologists.

[92] M. Tokunaga, N. Imamoto, and K. Sakata-Sogawa. Highly inclined thin illumination enables clear single-molecule imaging in cells. Nat. Methods, 5(2):159–161, 2008.

[93] P. Vallotton, A. Ponti, C. Waterman-Storer, E. Salmon, and G. Danuser. Recovery, visualization, and analysis of actin and tubulin polymer flow in live cells: A fluorescent speckle microscopy study. Biophys. J., 85(2):1289, 2003.

[94] R. van de Schoot, S. Depaoli, R. King, B. Kramer, K. Märtens, M. G. Tadesse, M. Vannucci, A. Gelman, D. Veen, J. Willemsen, et al. Bayesian statistics and modelling. Nat. Rev. Methods Primers, 1(1):1, 2021.

[95] F. J. Verweij, L. Balaj, C. M. Boulanger, D. R. Carter, E. B. Compeer, G. D’angelo, S. El Andaloussi, J. G. Goetz, J. C. Gross, V. Hyenne, et al. The power of imaging to understand extracellular vesicle biology in vivo. Nat. Methods, 18(9):1013, 2021.

[96] L. von Diezmann, Y. Shechtman, and W. Moerner. Three-dimensional localization of single molecules for super-resolution imaging and single-particle tracking. Chem. Rev., 117(11):7244, 2017.

[97] U. von Toussaint. Bayesian inference in physics. Rev. Mod. Phys., 83:943, 2011.

[98] L. Wasserman. All of nonparametric statistics. Springer Science & Business Media, 2006.

[99] N. P. Wells, G. A. Lessard, P. M. Goodwin, M. E. Phipps, P. J. Cutler, D. S. Lidke, B. S. Wilson, and J. H. Werner. Time-resolved three-dimensional molecular tracking in live cells. Nano Lett., 10(11):4732, 2010.

[100] N. P. Wells, G. A. Lessard, and J. H. Werner. Confocal, three-dimensional tracking of individual quantum dots in high-background environments. Anal. Chem., 80(24):9830, 2008.

[101] K. I. Willig, S. O. Rizzoli, V. Westphal, R. Jahn, and S. W. Hell. Sted microscopy reveals that synaptotagmin remains clustered after synaptic vesicle exocytosis. Nature, 440(7086):935, 2006.

[102] M. Winter, E. Wait, B. Roysam, S. K. Goderie, R. A. N. Ali, E. Kokovay, S. Temple, and A. R. Cohen. Vertebrate neural stem cell segmentation, tracking and lineaging with validation and editing. Nat. Protoc., 6(12):1942, 2011.

[103] M. R. Winter, C. Fang, G. Banker, B. Roysam, and A. R. Cohen. Axonal transport analysis using multitemporal association tracking. Int. J. Comput. Biol. Drug Des., 5(1):35, 2012.

[104] Q. Xue and M. C. Leake. A novel multiple particle tracking algorithm for noisy in vivo data by minimal path optimization within the spatio-temporal volume. In 2009 IEEE International Symposium on Biomedical Imaging: From Nano to Macro, page 1158, 2009.

[105] L. Yang, Z. Qiu, A. H. Greenaway, and W. Lu. A new framework for particle detection in low-snr fluorescence live-cell images and its application for improved particle tracking. IEEE Trans. Biomed. Eng., 59(7):2040, 2012.

[106] Z. Yin, T. Kanade, and M. Chen. Understanding the phase contrast optics to restore artifact-free microscopy images for segmentation. Med. Image Anal., 16(5):1047, 2012.

[107] B. Zhang, J. Zerubia, and J.-C. Olivo-Marin. Gaussian approximations of fluorescence microscope point-spread function models. Appl. Opt., 46(10):1819, 2007.

