## Supplementary Information for "BNP-Track: A framework for superresolved tracking"

**D Frame of reference** **18**

**E Time schedule** **19**

**F Optics representation** **20**

**M Definitions** **34**

**References** **35**

### A Additional figures

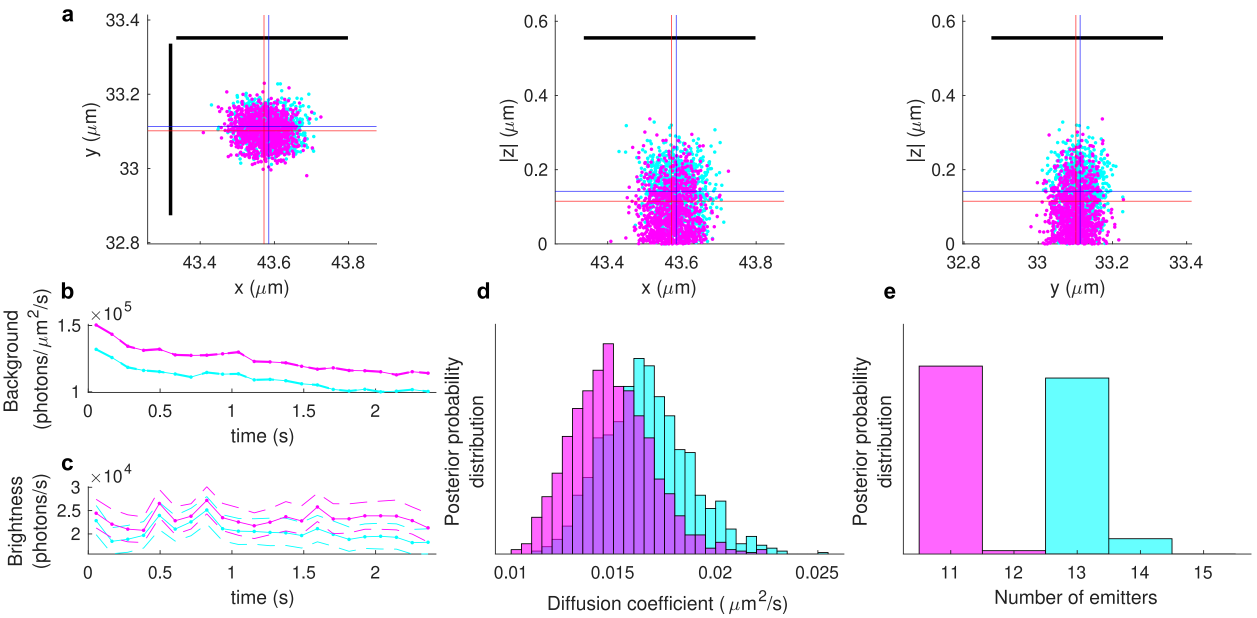

***Figure A.1:*** *Testing BNP-Track's performance for ROI-2 shown in fig. 2. The color scheme here is the same as in fig. 2 (cyan for camera A and magenta for camera B).* **a**, *Localization estimates in the lateral and axial directions at a selected frame of a selected emitter. Dots indicate individual positions sampled from the joint posterior distribution (as detailed in* METHODS*) and blue and red crosses indicate average values for cyan and magenta, respectively. The black line segments mark the diffraction limit in the lateral direction.* **b** *and* **c**, *Estimated background photon fluxes and emitter brightnesses for both cameras throughout the course of imaging. Dotted lines represent median estimates and dashed lines map the 1%-99% credible interval.* **d** *and* **e**, *The posterior distributions of diffusion coefficient and number of emitters for both cameras.*

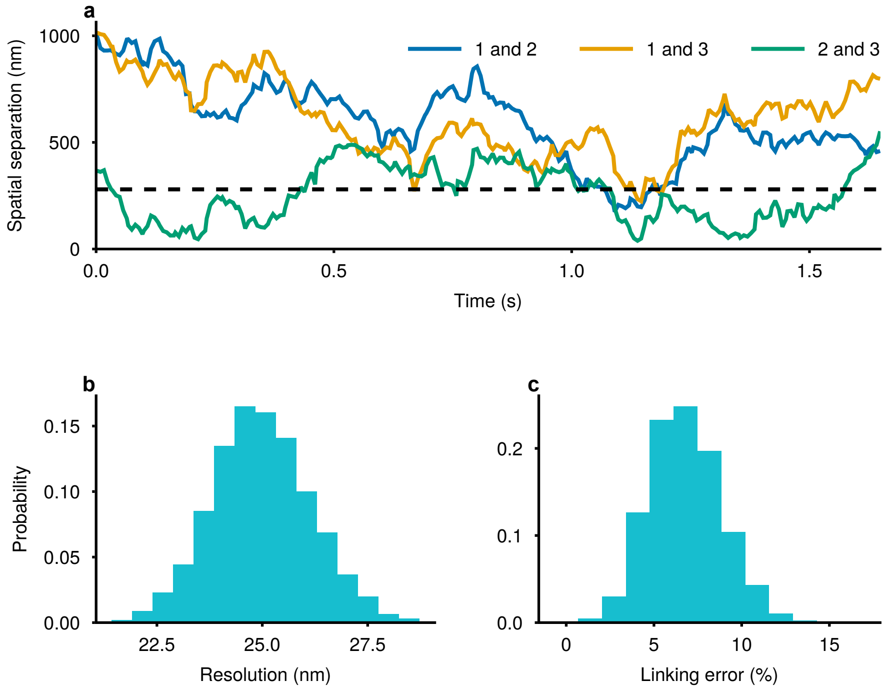

***Figure A.2:*** **a**, *The spatial separation between each emitter pair in Supplementary Video 7 as a function of time and the Rayleigh diffraction limit, is marked by the black dashed line.* **b**, *The distribution of BNP-Track's localization resolution distribution over all the sampled tracks. See the* RESULTS *section in main text for the definition of localization resolution.* **c**, *The distribution of incorrect linking percentages. Both histograms in this figure are normalized such that bin heights are probabilities.*

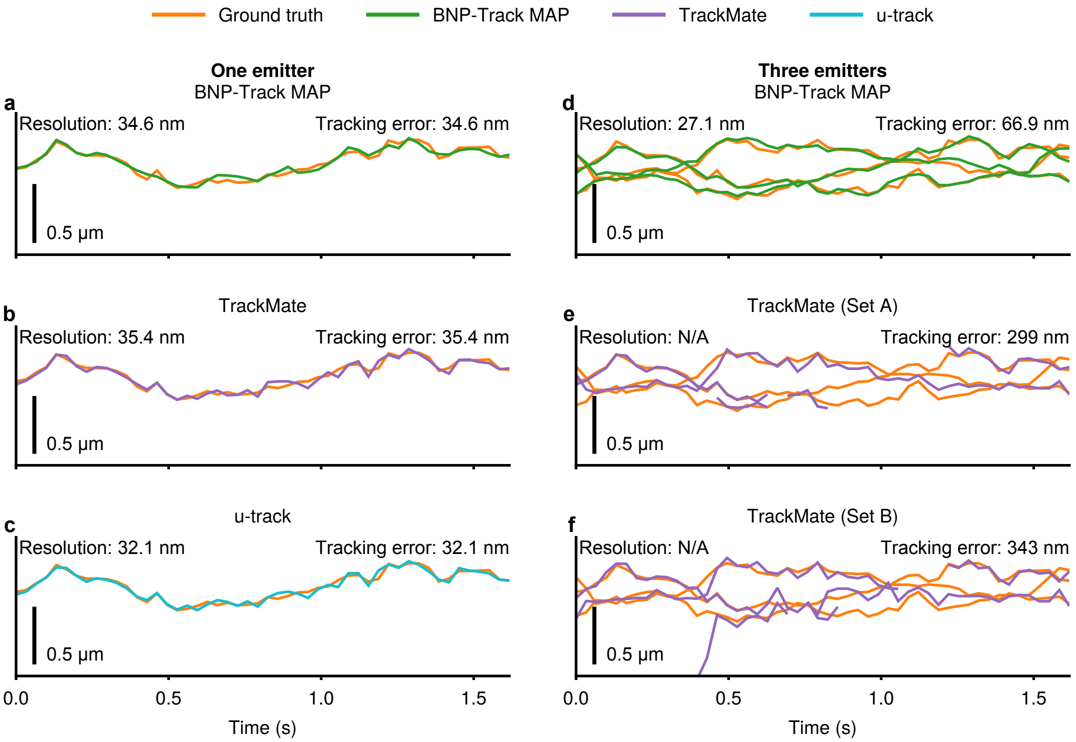

***Figure A.3:*** *Same layout as fig. 3 but for the $x$ coordinate.*

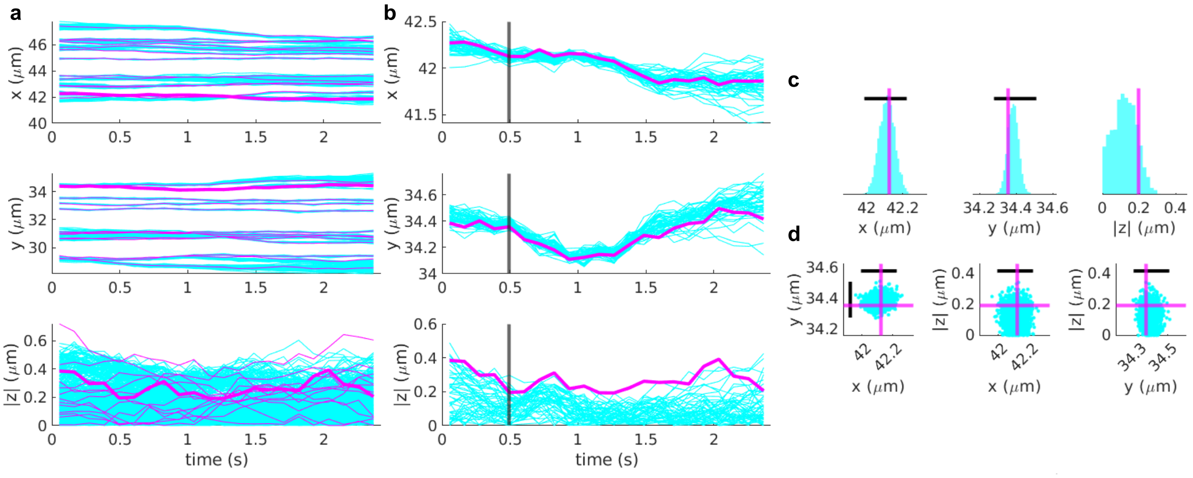

***Figure A.4:*** *BNP-Track's performance using synthetic data. Through all panels, BNP-Track's samples are in cyan and ground truths are in magenta.* **a**, *BNP-Track's reconstructions of time courses for all emitters' tracks.* **b**, *The track reconstructions of an out-of-focus emitter. This emitter's ground truth track is highlighted in* **a** *with thicker lines.* **c**, *Posterior distributions of individual localizations corresponding to the time level highlighted in* **b** *with black vertical lines. The black line segments mark the lateral diffraction limit in* **c**. **d**, *Sampled emitter locations in the lateral and axial directions. Again, crosses show estimate means and black line segments mark the lateral diffraction limit.*

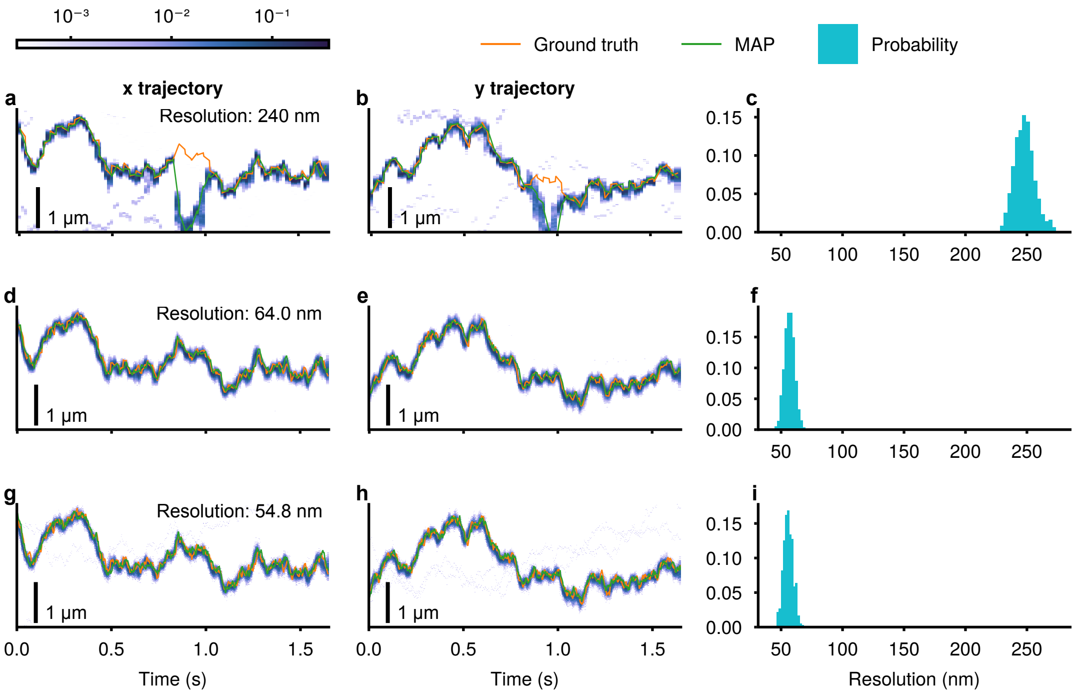

***Figure A.5:*** *BNP-Track's performance testing the number of positions sampled within the exposure period of one frame. This synthetic video (Supplementary Video 10) has one emitter diffusing at 1 $\mu m^2 s^{-1}$. In* **a**-**c** *BNP-Track only samples two positions within each exposure, while in* **d**-**f** *there are five, and in* **g**-**i** *there are ten. For this faster diffusing case, inferring two position per frame fails to correctly track the emitter in some frames, yielding a poor localization resolution, see (***c***). Once the number of positions sampled within one exposure is sufficient (greater than or equal to five in this case), increasing this number will no longer improve the resolution, see (***f***) and (***i***).*

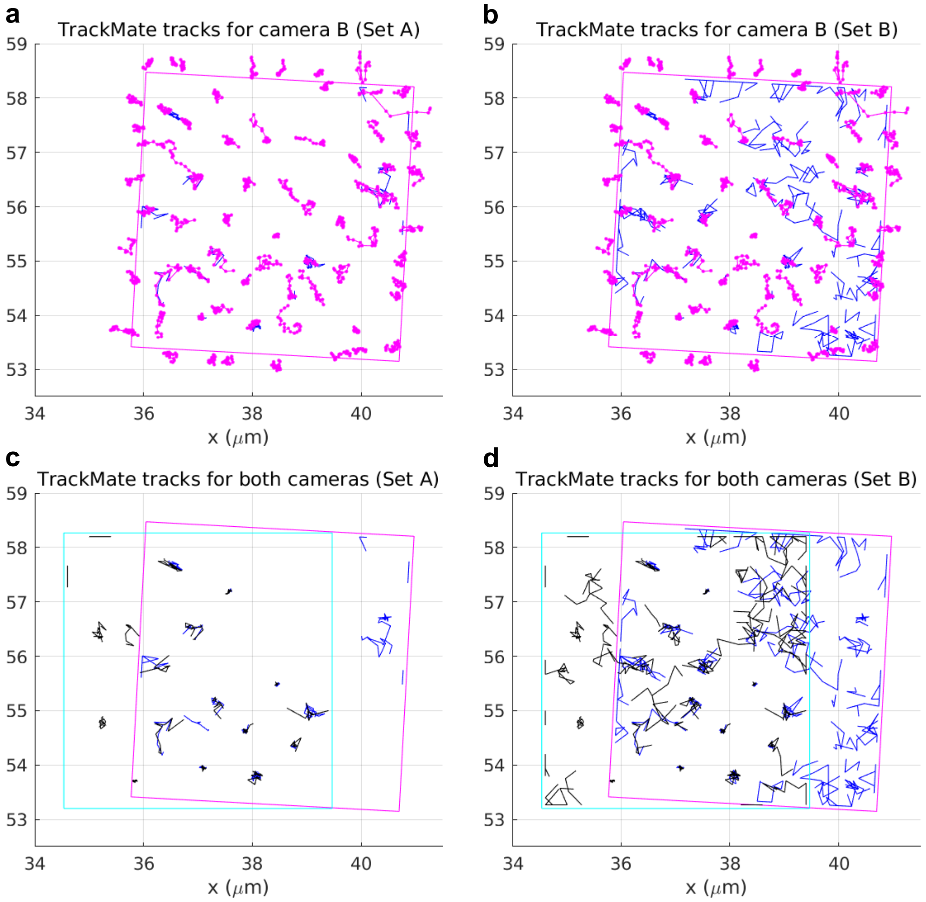

***Figure A.6:*** **a**, *A comparison between BNP-Track's result and TrackMate's result with a high localization quality threshold (Set A) for the data from camera B. TrackMate tracks are in blue.* **b**, *A comparison between BNP-Track's result and TrackMate's result with a low localization quality threshold (Set B) for the data from camera B. TrackMate tracks are in blue.* **c** *Comparison between TrackMate tracks (Set A) from both cameras with the same image registration as fig. 6.* **d** *Comparison between TrackMate tracks (Set B) from both cameras.*

#### B Additional tables

| Description | Parameter | Unit | Value |
|---|---|---|---|
| Emission wavelength | $\lambda$ | nm | 665 |
| Exposure period | $\tau$ | s | 0.03 |
| Frame period | | s | 0.033 |
| Frame height | | pixel | 23 |
| Frame with | | pixel | 15 |
| Numerical aperture | NA | | 1.45 |
| Pixel size | | nm | 133 |
| Refractive index | $n_{\mathsf{RI}}$ | | 1.515 |
| Noise excess factor | $f$ | | 2 |
| Quantum efficiency | $\beta$ | | |

***Table B.1:*** *List of parameter values used in data simulation.*

| | BNP-Track | TrackMate A | u-track |
|---|---|---|---|
| **Global measures** | | | |
| Pairing distance | 13.0 | 13.3 | 12.1 |
| Normalized pairing score (alpha) | 0.948 | 0.947 | 0.952 |
| Full normalized score (beta) | 0.948 | 0.947 | 0.952 |
| **Tracks** | | | |
| Number of reference tracks | 1 | 1 | 1 |
| Number of candidate tracks | 1 | 1 | 1 |
| Similarity between tracks (Jaccard) | 1.0 | 1.0 | 1.0 |
| Number of paired tracks | 1 | 1 | 1 |
| Number of missed tracks | 0 | 0 | 0 |
| Number of spurious tracks | 0 | 0 | 0 |
| **Detections** | | | |
| Number of reference detections | 50 | 50 | 50 |
| Number of candidate detections | 50 | 50 | 50 |
| Similarity between detections (Jaccard) | 1.0 | 1.0 | 1.0 |
| Number of paired detections | 50 | 50 | 50 |
| Number of missed detections | 0 | 0 | 0 |
| Number of spurious detections | 0 | 0 | 0 |
| **Detection accuracy** | | | |
| Root mean-square error | 0.300 | 0.314 | 0.297 |
| Minimum distance | 0.0310 | 0.0168 | 0.0241 |
| Maximum distance | 0.660 | 0.809 | 0.888 |
| Distance standard deviation | 0.150 | 0.166 | 0.173 |

***Table B.2:*** *Full tracking performance measure comparison among BNP-Track, TrackMate, and u-track for the one-emitter video (Supplementary Video 6).*

| | BNP-Track | TrackMate A | TrackMate B |
|---|---|---|---|
| Global measures | | | |
| Pairing distance | 75.4 | 337 | 386 |
| Normalized pairing score (alpha) | 0.899 | 0.550 | 0.484 |
| Full normalized score (beta) | 0.899 | 0.550 | 0.484 |
| Tracks | | | |
| Number of reference tracks | 3 | 3 | 3 |
| Number of candidate tracks | 3 | 3 | 3 |
| Similarity between tracks (Jaccard) | 1 | 1 | 1 |
| Number of paired tracks | 3 | 3 | 3 |
| Number of missed tracks | 0 | 0 | 0 |
| Number of spurious tracks | 0 | 0 | 0 |
| Detections | | | |
| Number of reference detections | 150 | 150 | 150 |
| Number of candidate detections | 150 | 106 | 93 |
| Similarity between detections (Jaccard) | 1.0 | 0.593 | 0.62 |
| Number of paired detections | 150 | 89 | 93 |
| Number of missed detections | 0 | 61 | 57 |
| Number of spurious detections | 0 | 0 | 1 |
| Detection accuracy | | | |
| Root mean-square error | 0.807 | 0.473 | 1.77 |
| Minimum distance | 0.0441 | 0.0242 | 0.0227 |
| Maximum distance | 3.62 | 1.07 | 4.65 |
| Distance standard deviation | 0.631 | 0.303 | 1.40 |

***Table B.3:*** *Full tracking performance measure comparison between BNP-Track and TrackMate for the three-emitter video (Supplementary Video 7).*

| | BNP-Track | TrackMate |
|---|---|---|
| Global measures | | |
| Pairing distance (pixel) | 10.9 | 74.1 |
| Normalized pairing score (alpha) | 0.950 | 0.663 |
| Full normalized score (beta) | 0.950 | 0.663 |
| Tracks | | |
| Number of reference tracks | 2 | 2 |
| Number of candidate tracks | 2 | 2 |
| Similarity between tracks (Jaccard) | 1.0 | 1.0 |
| Number of paired tracks | 2 | 2 |
| Number of missed tracks | 0 | 0 |
| Number of spurious tracks | 0 | 0 |
| Detections | | |
| Number of reference detections | 44 | 44 |
| Number of candidate detections | 44 | 38 |
| Similarity between detections (Jaccard) | 1.0 | 0.795 |
| Number of paired detections | 44 | 35 |
| Number of missed detections | 0 | 9 |
| Number of spurious detections | 0 | 0 |
| Detection accuracy in pixel | | |
| Root mean-square error | 0.277 | 1.13 |
| Minimum distance | 0.0275 | 0.101 |
| Maximum distance | 0.511 | 3.54 |
| Distance standard deviation | 0.122 | 0.767 |
| Detection accuracy in nm | | |
| Root mean-square error | 36.8 | 150 |
| Minimum distance | 3.66 | 13.4 |
| Maximum distance | 68.0 | 471 |
| Distance standard deviation | 16.2 | 102 |

***Table B.4:*** *Full tracking performance measure comparison between BNP-Track and TrackMate for the synthetic video in fig. 5a, 5g, and 5m.*

| | BNP-Track | TrackMate |
|---|---|---|
| Global measures | | |
| Pairing distance (pixel) | 8.67 | 110 |
| Normalized pairing score (alpha) | 0.961 | 0.501 |
| Full normalized score (beta) | 0.961 | 0.501 |
| Tracks | | |
| Number of reference tracks | 2 | 2 |
| Number of candidate tracks | 2 | 2 |
| Similarity between tracks (Jaccard) | 1.0 | 1.0 |
| Number of paired tracks | 2 | 2 |
| Number of missed tracks | 0 | 0 |
| Number of spurious tracks | 0 | 0 |
| Detections | | |
| Number of reference detections | 44 | 44 |
| Number of candidate detections | 44 | 31 |
| Similarity between detections (Jaccard) | 1.0 | 0.659 |
| Number of paired detections | 44 | 29 |
| Number of missed detections | 0 | 15 |
| Number of spurious detections | 0 | 0 |
| Detection accuracy in pixel | | |
| Root mean-square error | 0.221 | 1.64 |
| Minimum distance | 0.0439 | 0.103 |
| Maximum distance | 0.495 | 3.73 |
| Distance standard deviation | 0.101 | 1.12 |
| Detection accuracy in nm | | |
| Root mean-square error | 29.4 | 218 |
| Minimum distance | 5.84 | 13.7 |
| Maximum distance | 65.8 | 496 |
| Distance standard deviation | 13.4 | 149 |

***Table B.5:*** *Full tracking performance measure comparison between BNP-Track and TrackMate for the synthetic video in fig. 5b, 5h, and 5n.*

| | BNP-Track | TrackMate |
|---|---|---|
| Global measures | | |
| Pairing distance (pixel) | 12.1 | 111 |
| Normalized pairing score (alpha) | 0.945 | 0.498 |
| Full normalized score (beta) | 0.945 | 0.498 |
| Tracks | | |
| Number of reference tracks | 2 | 2 |
| Number of candidate tracks | 2 | 2 |
| Similarity between tracks (Jaccard) | 1.0 | 1.0 |
| Number of paired tracks | 2 | 2 |
| Number of missed tracks | 0 | 0 |
| Number of spurious tracks | 0 | 0 |
| Detections | | |
| Number of reference detections | 44 | 44 |
| Number of candidate detections | 44 | 27 |
| Similarity between detections (Jaccard) | 1.0 | 0.614 |
| Number of paired detections | 44 | 27 |
| Number of missed detections | 0 | 17 |
| Number of spurious detections | 0 | 0 |
| Detection accuracy in pixel | | |
| Root mean-square error | 0.302 | 1.05 |
| Minimum distance | 0.0353 | 0.290 |
| Maximum distance | 0.508 | 2.09 |
| Distance standard deviation | 0.125 | 0.454 |
| Detection accuracy in nm | | |
| Root mean-square error | 40.2 | 140 |
| Minimum distance | 4.69 | 38.6 |
| Maximum distance | 67.6 | 278 |
| Distance standard deviation | 16.6 | 60.4 |

***Table B.6:*** *Full tracking performance measure comparison between BNP-Track and TrackMate for the synthetic video in fig. 5c, 5i, and 5o.*

| | BNP-Track | TrackMate |
|---|---|---|
| Global measures | | |
| Pairing distance (pixel) | 12.2 | 122 |
| Normalized pairing score (alpha) | 0.945 | 0.443 |
| Full normalized score (beta) | 0.945 | 0.443 |
| Tracks | | |
| Number of reference tracks | 2 | 2 |
| Number of candidate tracks | 2 | 1 |
| Similarity between tracks (Jaccard) | 1.0 | 0.5 |
| Number of paired tracks | 2 | 1 |
| Number of missed tracks | 0 | 1 |
| Number of spurious tracks | 0 | 0 |
| Detections | | |
| Number of reference detections | 44 | 44 |
| Number of candidate detections | 44 | 22 |
| Similarity between detections (Jaccard) | 1.0 | 0.5 |
| Number of paired detections | 44 | 22 |
| Number of missed detections | 0 | 22 |
| Number of spurious detections | 0 | 0 |
| Detection accuracy in pixel | | |
| Root mean-square error | 0.309 | 0.694 |
| Minimum distance | 0.0195 | 0.0366 |
| Maximum distance | 0.664 | 1.91 |
| Distance standard deviation | 0.136 | 0.401 |
| Detection accuracy in nm | | |
| Root mean-square error | 41.1 | 92.3 |
| Minimum distance | 2.59 | 4.87 |
| Maximum distance | 88.3 | 254 |
| Distance standard deviation | 18.1 | 53.3 |

***Table B.7:*** *Full tracking performance measure comparison between BNP-Track and TrackMate for the synthetic video in fig. 5d, 5j, and 5p.*

| | BNP-Track | TrackMate |
|---|---|---|
| Global measures | | |
| Pairing distance (pixel) | 12.0 | 118 |
| Normalized pairing score (alpha) | 0.946 | 0.464 |
| Full normalized score (beta) | 0.946 | 0.464 |
| Tracks | | |
| Number of reference tracks | 2 | 2 |
| Number of candidate tracks | 2 | 2 |
| Similarity between tracks (Jaccard) | 1.0 | 1.0 |
| Number of paired tracks | 2 | 2 |
| Number of missed tracks | 0 | 0 |
| Number of spurious tracks | 0 | 0 |
| Detections | | |
| Number of reference detections | 44 | 44 |
| Number of candidate detections | 44 | 23 |
| Similarity between detections (Jaccard) | 1.0 | 0.523 |
| Number of paired detections | 44 | 23 |
| Number of missed detections | 0 | 21 |
| Number of spurious detections | 0 | 0 |
| Detection accuracy in pixel | | |
| Root mean-square error | 0.300 | 0.633 |
| Minimum distance | 0.0599 | 0.0254 |
| Maximum distance | 0.566 | 1.15 |
| Distance standard deviation | 0.127 | 0.293 |
| Detection accuracy in nm | | |
| Root mean-square error | 39.9 | 84.2 |
| Minimum distance | 7.97 | 3.38 |
| Maximum distance | 75.3 | 153 |
| Distance standard deviation | 16.9 | 39.0 |

***Table B.8:*** *Full tracking performance measure comparison between BNP-Track and TrackMate for the synthetic video in fig. 5e, 5k, and 5q.*

| | BNP-Track | TrackMate |
|---|---|---|
| Global measures | | |
| Pairing distance (pixel) | 16.47 | 121 |
| Normalized pairing score (alpha) | 0.925 | 0.452 |
| Full normalized score (beta) | 0.925 | 0.452 |
| Tracks | | |
| Number of reference tracks | 2 | 2 |
| Number of candidate tracks | 2 | 1 |
| Similarity between tracks (Jaccard) | 1.0 | 0.5 |
| Number of paired tracks | 2 | 1 |
| Number of missed tracks | 0 | 1 |
| Number of spurious tracks | 0 | 0 |
| Detections | | |
| Number of reference detections | 44 | 44 |
| Number of candidate detections | 44 | 22 |
| Similarity between detections (Jaccard) | 1.0 | 0.5 |
| Number of paired detections | 44 | 22 |
| Number of missed detections | 0 | 22 |
| Number of spurious detections | 0 | 0 |
| Detection accuracy in pixel | | |
| Root mean-square error | 0.420 | 0.576 |
| Minimum distance | 0.0803 | 0.0620 |
| Maximum distance | 1.09 | 1.16 |
| Distance standard deviation | 0.191 | 0.321 |
| Detection accuracy in nm | | |
| Root mean-square error | 55.9 | 76.6 |
| Minimum distance | 10.7 | 8.25 |
| Maximum distance | 145 | 154 |
| Distance standard deviation | 25.4 | 42.7 |

***Table B.9:*** *Full tracking performance measure comparison between BNP-Track and TrackMate for the synthetic video in fig. 5f, 5l, and 5r.*

### C Probabilistic modeling

Because most of the notions in this study are stochastic, we use probabilistic relations and statistical notations. The methodology we adopt facilitates the description of the variables in BNP-Track and the relations among them. Since the statistical conventions we follow are standard, here we provide only a brief description. For a complete presentation, we refer to [9, 53, 59, 18]. Also, a comprehensive introduction to the computational schemes we implement can be found in [59, 48] and complete descriptions in [18, 47, 35].

#### C.1 Statistical notation

A statistical notation like $\theta \sim \mathrm{P}$ indicates that $\theta$ is a random variable and this random variable follows the probability distribution P. This means that in our computations the values of $\theta$ follow the probability density $p(\theta)$ that is associated with P. For example, $\theta \sim \mathrm{Normal}\left(\mu, \sigma^2\right)$, which means that $\theta$ is a normal random variable with mean $\mu$ and variance $\sigma^2$, indicates that $\theta$ is distributed according to the density $p(\theta) = \frac{1}{\sqrt{2\pi\sigma^2}} \exp\left(-\frac{(\mu-\theta)^2}{2\sigma^2}\right)$.

Further, a notation like $w|\theta \sim \mathrm{P}_\theta$ indicates that $w$ is a random variable whose distribution depends on another random variable $\theta$. For example, $w|\theta \sim \mathrm{Normal}\left(\theta, s^2\right)$, indicates that $w$ is distributed according to the density $p(w|\theta) = \frac{1}{\sqrt{2\pi s^2}} \exp\left(-\frac{(\theta-w)^2}{2s^2}\right)$.

Our random variables do not necessarily have to be univariate. As such, $w$ or $\theta$ can encompass multiple individual random variables that describe various aspects of our image processing model. The densities $p(w)$ and $p(w|\theta)$ may also depend on other non-random parameters, which we do not explicitly include on the left-hand side of statistical equations.

We provide a summary of the distributions we use in this study and their densities in table M.1. Because some distributions do not have a standardized parameterization, this table clarifies also our particular choices.

#### C.2 Statistical inference

The probability density of two random variables, $w$ and $\theta$, denoted by $p(w, \theta)$, can be expressed as the product of their individual density functions, i.e., $p(w, \theta) = p(w|\theta)p(\theta)$. This relationship is symmetric, as shown by the equality $p(w, \theta) = p(\theta, w) = p(w|\theta)p(\theta) = p(\theta|w)p(w)$. Therefore, the density function $p(\theta|w)$ is proportional to the product $p(w|\theta)p(\theta)$. In other words, the specification of the distributions of $\theta \sim \mathrm{P}$ and $w|\theta \sim \mathrm{P}_\theta$ is sufficient to derive the density function $p(\theta|w)$, and no other information is required.

Throughout this study, we use $w$ to gather the image measurements and $\theta$ to gather the variables whose values we seek to estimate. In this setting, the distributions of $\theta$ and $\theta|w$, or their densities $p(\theta)$ and $p(\theta|w)$, are designated as prior and posterior, respectively. These two are linked via $w|\theta$, or its density $p(w|\theta)$, which is designated as likelihood. As we demonstrate in RESULTS, our approach focuses on the posterior $p(\theta|w)$. In METHODS, we describe this posterior by formulating each factor $p(w|\theta)$ and $p(\theta)$ separately. On appendices H and I we describe in detail how each of these factors is represented mathematically.

#### C.3 Statistical simulation

While the posterior distribution $p(\theta|w)$ is proportional to the product of $p(w|\theta)$ and $p(\theta)$, there is no mathematical formula that enables direct evaluation of this distribution. However, in this paper, we present a method for computing the posterior indirectly through sampling simulations. Specifically, we generate a sequence of values

$\theta^{(1)}, \theta^{(2)}, \theta^{(3)}, \ldots$ that have the same statistical properties as $p(\theta|w)$. We use this sequence to produce results such as histograms, scatter-plots, mean values, and credible intervals without relying on an analytic formula for $p(\theta|w)$. In appendix K, we provide a detailed description of how to efficiently carry out posterior sampling in BNP-Track.

To facilitate the interpretation of our results, we present a simplified example in fig. C.1 where we simulate from a toy posterior $p(\theta|w)$. In this example, we consider a bivariate random variable $\theta = (\theta_1, \theta_2)$. Therefore, we generate two simultaneous sequences of samples, $\theta_1^{(1)}, \theta_1^{(2)}, \theta_1^{(3)}, \ldots$ and $\theta_2^{(1)}, \theta_2^{(2)}, \theta_2^{(3)}, \ldots$, with the specific values shown in the left panels. Using these sequences, we construct histograms in the middle panels to visualize the distribution of each variable, and a scatter plot in the right panel to illustrate the correlation between $\theta_1$ and $\theta_2$. It is important to note that if our posterior consisted of more than two random variables, such as the posterior described in METHODS, we would require more than two simultaneous sequences of samples to generate more than two histograms and multiple scatter plots to fully visualize all correlations developed.

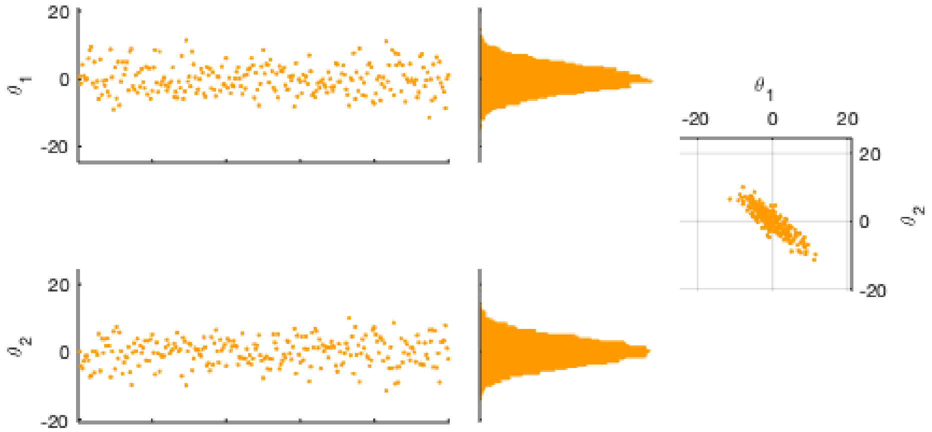

***Figure C.1:*** *Sampling from a toy posterior $p(\theta|w)$ with a bivariate variable $\theta = (\theta_1, \theta_2)$ fully characterizes the statistics of the involved variables as well as the correlations among them.*

### D Frame of reference

In this study, we exclusively consider an object space frame of reference. Specifically, in the object space we consider Cartesian coordinates $x, y, z$ as illustrated on the left side of fig. D.1. Consistent with common practice [20, 16, 52, 10], our $z$ axis is parallel to the optical axis of the microscope and points away from the image plane. Specifically, on an inverted microscope, the $z$ axis points upwards. Moreover, we assume that the microscope and the camera are aligned such that the image plane is perpendicular to the optical axis [20, 16, 52, 10]. Lastly, we orient the $x, y$ axes in such a way that they are parallel to the pixel edges of the camera. Overall, this coordinate system follows the right-hand rule.

Under these conventions, the image plane, physically lying on the camera, can be projected to the object plane, physically located on the object space. The object plane is the plane in the sample space yielding focused images, as shown on the right of fig. D.1. Assuming that the origins in the object space and image plane are conjugated, image plane coordinates $x', y'$ are related to object space coordinates $x, y, z$ by

$$x = \frac{x'}{\mathcal{M}}, \qquad y = \frac{y'}{\mathcal{M}}, \qquad z = 0$$

where $\mathcal{M}$ is the combined magnification achieved through the microscope's objective and tube lenses. With this convention, a square pixel with physical size of 16 $\mu$m, on a microscope imaging at 120x, is projected on a pixel of size $133.33$ nm.

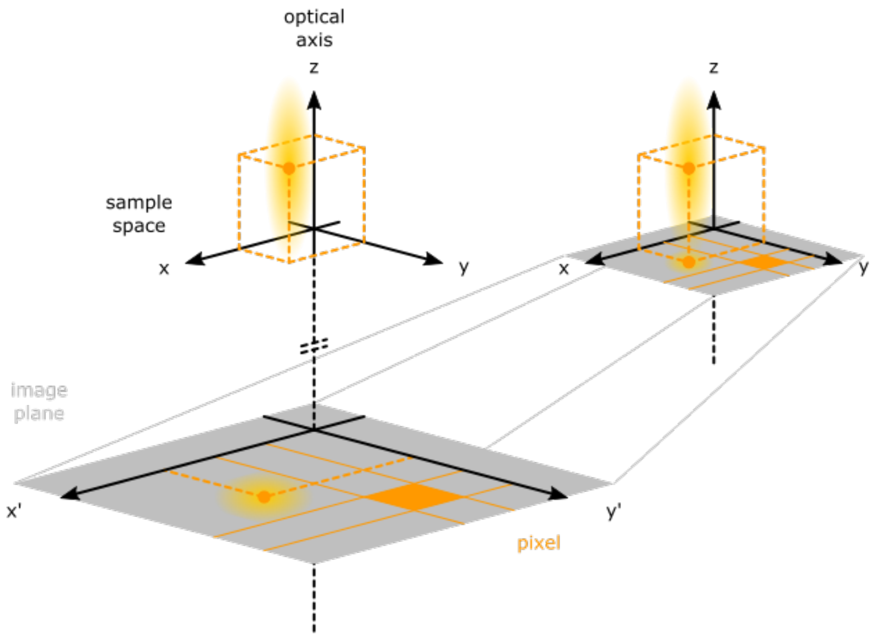

***Figure D.1:*** *Frames of reference. Upper left: object space, located in front of the microscope's lenses along the optical path. Lower left: image plane, located behind the microscope's lenses along the optical path. Right: object space with the image plane projected on the object plane. In this study, we use exclusively the latter configuration where the image plane is identified with the object plane and position coordinates are expressed in sample space units.*

### E Time schedule

In this study, we refer to time in relation to the acquisition schedule of the images, which is determined by the camera used. The acquisition of each image involves two phases: an integration period, during which the camera pixels are sensitive to incident photons, and a dead period, during which the camera pixels remain unresponsive to incident photons [32, 33, 50]. To improve the accuracy of dynamical approximations, as we describe in detail in appendix G, we model the dead time associated with image acquisition as occurring in two equal periods: one immediately before and one immediately after the corresponding integration period. Our convention is illustrated in fig. E.1. Analytically, our temporal frame of reference is as follows.

We label the images sequentially with $n = 1, \dots, N$, where $n = 1$ marks the earliest acquired image in the experiment session and $n = N$ the latest. As shown in fig. E.1, we denote with $t_n^{\text{min}}$ and $t_n^{\text{max}}$ the start and end times of the integration period of the $n^{\text{th}}$ image. We report our estimates at times, which we denote with $t_n$ and label with $n = 0, 1, \dots, N$, that separate successive image acquisitions. Specifically, we distribute the total dead time $\tau_n^{\text{dead}}$, associated with the $n^{\text{th}}$ image, equally between the time intervals immediately preceding $t_n^{\text{min}}$ and following $t_n^{\text{max}}$. With this choice, the acquisition of the $n^{\text{th}}$ image is separated by the preceding and following one at the respective times $t_{n-1} = t_n^{\text{min}} - \tau_n^{\text{dead}}/2$ and $t_n = t_n^{\text{max}} + \tau_n^{\text{dead}}/2$.

Provided the frame rate $\nu^{\text{frame}}$ and dead time $\tau^{\text{dead}}$ remain constant throughout the imaging course, as it is common in most experiments, our convention becomes

$$t_n = \frac{n}{\nu^{\text{frame}}}, \qquad t_n^{\text{min}} = t_{n-1} + \frac{\tau^{\text{dead}}}{2}, \qquad t_n^{\text{max}} = t_n - \frac{\tau^{\text{dead}}}{2}.$$

For example, with this convention, 5 images acquired at a frame rate of 10 Hz and a dead time of 10 ms, correspond to

$$t_1 = 100 \text{ ms}, \qquad t_2 = 200 \text{ ms}, \qquad t_3 = 300 \text{ ms}, \qquad t_4 = 400 \text{ ms}, \qquad t_5 = 500 \text{ ms},$$

and the individual integration periods last between

$$\begin{aligned} t_1^{\text{min}} &= 5 \text{ ms}, & t_2^{\text{min}} &= 105 \text{ ms}, & t_3^{\text{min}} &= 205 \text{ ms}, & t_4^{\text{min}} &= 305 \text{ ms}, & t_5^{\text{min}} &= 405 \text{ ms}, \\ t_1^{\text{max}} &= 95 \text{ ms}, & t_2^{\text{max}} &= 195 \text{ ms}, & t_3^{\text{max}} &= 295 \text{ ms}, & t_4^{\text{max}} &= 395 \text{ ms}, & t_5^{\text{max}} &= 495 \text{ ms}. \end{aligned}$$

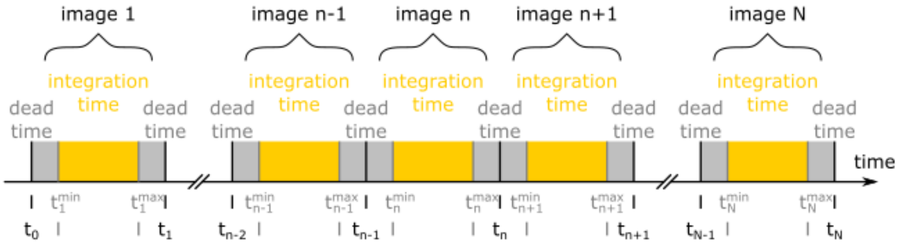

***Figure E.1:*** *Temporal frame of reference. In our convention, the acquisition of each image contains three separate phases: a dead time phase right before the integration time; an integration time phase; and a dead time phase right after the integration time. Specifically, for the $n^{th}$ image the three phases are distinguished by the times $t_{n-1}, t_n^{\text{min}}, t_n^{\text{max}}, t_n$.*

#### F Optics representation

Due to the incoherent imaging conditions encountered in fluorescence microscopy [20, 16, 52, 10, 33], we describe the optical response of the imaging apparatus with an additive photon flux of the form

$$U(x,y,t) = U_{\text{back}}(x,y,t) + \sum_{m=1}^{M} b^m U^m_{\text{fluor}}(x,y,t)$$

that sums up background photon contributions $U_{\text{back}}(x,y,t)$ and emitter photon contributions $U^m_{\text{fluor}}(x,y,t)$. Our photon flux $U(x,y,t)$ models the irradiance [50, 60, 58] on the image plane, which according to appendix D we project to the object plane in the sample space, as it changes over the time course of the imaging experiment $t$.

In turn, we model $U_{\mathsf{back}}(x,y,t) = C(t)$ as a uniform-in-space flux, which we denote with $C(t)$; while, we model fluorophore contributions by summing flux contributions of each emitter $U^m_{\text{fluor}}(x,y,t)$ which we label with $m = 1,\dots,M$. We model the latter by the product

$$U^m_{\text{fluor}}(x,y,t) = h(t)\, G_g\left(x,y; X^m(t), Y^m(t), Z^m(t)\right).$$

This product consists of a point spread function $G_g(x,y;X,Y,Z)$, which models how the incident photons stemming from a light emitter at position $X,Y,Z$ are spread over the image plane [2, 20, 16, 10, 19, 55]; and, a photon emission rate $h(t)$, which describes how often photons incident on the image plane are emitted by a single light emitter.

From the two factors forming each $U^m_{\text{fluor}}(x,y,t)$, the point spread function $G_g(x,y;X,Y,Z)$ is measured in units of 1/area and the photon emission rate $h(t)$ is measured in units of photons/time. Considered together, these result in a flux with units of photons/area/time.

In superresolution microscopy, the point spread function is primarily affected by diffraction and microscope specific aberrations [2, 34, 20, 16, 10, 19, 55] and the photon emission rate is set by the labeling fluorophores and the illumination modality applied [33, 50]. For these reasons, we obtain $G_g(x,y;X,Y,Z)$ by fitting calibration measurements, as described in appendix L, and $h(t)$ by fitting during processing, as described in appendix K.

##### F.1 Point spread function

For all light emitters, we consider Gaussian point spread functions [13, 20, 2, 61]. These have the form

$$G_g(x,y;X,Y,Z) = \frac{1}{2\pi\sigma_g^2(Z)} \exp\left(-\frac{1}{2}\frac{(x-X)^2 + (y-Y)^2}{\sigma_g^2(Z)}\right).$$

By convention, we normalize each point spread function to unit total volume, i.e. $\iint_{-\infty,-\infty}^{+\infty,+\infty} dxdy\, G_g(x,y;X,Y,Z) = 1$, so we can readily compare photon emissions from in-focus and out-of-focus emitters.

The extent of the point spread function, which is proportional to $\sigma_g(Z)$, depends upon the position $Z$ of the emitter along the optical axis [34, 10, 19, 55]. Assuming, the axial light propagation is similar to a Gaussian beam [13, 52], we model $\sigma_g(Z)$ by

$$\sigma_g(Z) = \sigma_{\text{ref}} \sqrt{g_1 + g_2 \left(\frac{Z}{Z_{\text{ref}}}\right)^2},$$

where $\sigma_{\text{ref}}$ and $Z_{\text{ref}}$ are reference values for our point spread function with units of length and $g_1, g_2$ are unitless parameters that allow our point spread function to accommodate microscope specific aberrations in the width of the focused point spread function and the depth of focus. Namely, $g_1$ adjusts for deviations in the width of the point spread function of an in-focus emitter, $Z = 0$; while, $g_2$ adjusts for deviations in the width of the point spread function of an out-of-focus emitter, $Z \neq 0$.

We obtain reference values $\sigma_{\text{ref}}, Z_{\text{ref}}$ considering diffraction limited imaging under ideal conditions. Specifically, following [2, 61, 10, 20, 50, 58] we use

$$\sigma_{\text{ref}} = 0.21 \frac{\lambda}{n_{\text{NA}}} \qquad\qquad Z_{\text{ref}} = 4\pi n_{\text{RI}} \frac{\sigma_{\text{ref}}^2}{\lambda}$$

where $n_{\text{NA}}$ denotes the microscope's numerical aperture, $\lambda$ denotes the imaged wavelength (in vacuum), and $n_{\text{RI}}$ denotes the index of refraction of the objective's immersion fluid. However, the values of $g_1, g_2$ need to be adjusted to fit each particular microscope. This is achieved following the calibration protocol in appendix L.

##### F.2 Effective pixel function

Combining the flux $U(x, y, t)$ with the pixel response, which integrates photon contributions over space and time [50, 33, 5], the average number of incident photons on the $p^{\text{th}}$ pixel during the $n^{\text{th}}$ exposure takes the form

$$\begin{aligned} u_n^p &= \int_{t_n^{\min}}^{t_n^{\max}} dt \iint_{x_{\min}^p, y_{\min}^p}^{x_{\max}^p, y_{\max}^p} dx dy \, U(x, y, t) \\ &= \int_{t_n^{\min}}^{t_n^{\max}} dt \left( C(t) A^p + h(t) \sum_{m=1}^{M} b^m \iint_{x_{\min}^p, y_{\min}^p}^{x_{\max}^p, y_{\max}^p} dx dy \, G_g(x, y; X^m(t), Y^m(t), Z^m(t)) \right) \end{aligned}$$

where $A^p = (x_{\max}^p - x_{\min}^p)(y_{\max}^p - y_{\min}^p)$ is the area monitored by the $p^{\text{th}}$ pixel.

To simplify the notation from now on, for each pixel, we consider an effective function, defined by

$$Q_g^p(X, Y, Z) = \iint_{x_{\min}^p, y_{\min}^p}^{x_{\max}^p, y_{\max}^p} dx dy \; G_g(x, y; X, Y, Z),$$

which combines the effects of diffraction and finite pixel size. Our effective functions are unitless. With this convention, the average number of photons incident on the $p^{\text{th}}$ pixel during the $n^{\text{th}}$ exposure becomes

$$u_n^p = \int_{t_n^{\min}}^{t_n^{\max}} dt \left( C(t) A^p + h(t) \sum_{m=1}^{M} b^m Q_g^p(X^m(t), Y^m(t), Z^m(t)) \right).$$

### G Dynamics representation

#### G.1 Temporal discretization

To proceed with image analysis, the average numbers of incident photons $u_{1:N}^{1:P}$ over all pixels and exposures need to be related to dynamical variables that are temporally discretized. For this, we approximate the integral of

$$u_n^p = \int_{t_n^{\min}}^{t_n^{\max}} dt \left( C(t)A^p + h(t) \sum_{m=1}^{M} b^m Q_g^p(X^m(t), Y^m(t), Z^m(t)) \right).$$

Analytically, our approximations rely on the following numerical quadrature formulas

$$\int_{t_n^{\min}}^{t_n^{\max}} dt\, C(t) \approx \tau_n^{\text{exps}} C_n,$$

$$\int_{t_n^{\min}}^{t_n^{\max}} dt\, h(t) Q_g^p(X^m(t), Y^m(t), Z^m(t)) \approx \tau_n^{\text{exps}} h_n Q_g^p(X_n^m, Y_n^m, Z_n^m),$$

where $\tau_n^{\text{exps}} = t_n^{\max} - t_n^{\min}$ is the integration time of the $n^{\text{th}}$ exposure and the discretized variables $C_{1:N}$, $h_{1:N}$, $X_{1:N}^{1:M}$, $Y_{1:N}^{1:M}$, and $Z_{1:N}^{1:M}$ are related to their continuous counterparts $C(t)$, $h(t)$, $X^{1:M}(t)$, $Y^{1:M}(t)$, and $Z^{1:M}(t)$ according to

$$C_n = C(T_n), \qquad h_n = h(T_n), \qquad X_n^m = X^m(T_n), \qquad Y_n^m = Y^m(T_n), \qquad Z_n^m = Z^m(T_n).$$

Finally, the discretized time levels $T_{1:N}$ are given by

$$T_n = \frac{t_n^{\min} + t_n^{\max}}{2}.$$

In essence, we use the mid-point rules [22, 51, 4, 46] to approximate the quadrature of $C(t)$, $h(t)$, $X^{1:M}(t)$, $Y^{1:M}(t)$, and $Z^{1:M}(t)$. As explained in appendix E, since the times $t_{0:N}$ are centered, both rules result in errors that decrease super-linearly with respect to the frame rate when the frame rate remains constant over the imaging course.

Ignoring the errors introduced by the quadrature formulas, from now on we consider average numbers of incident photons $u_{1:N}^{1:P}$ that we evaluate according to

$$u_n^p = \tau_n^{\text{exps}} \left( C_n A^p + h_n \sum_{m=1}^{M} b^m Q_g^p(X_n^m, Y_n^m, Z_n^m) \right).$$

Since in typical experiments background arises from uncharacterized sources and the photon emission rate is affected by uncontrolled factors, in this study, we do not model the variables $C_{1:N}$ and $h_{1:N}$ any further. However, to link each emitter's discretized positions $X_{1:N}^m$, $Y_{1:N}^m$, and $Z_{1:N}^m$ across time, we invoke physically realistic dynamical models of motion that we describe below.

#### G.2 Simplification for time-independent emitter brightness and background photon flux

As mentioned in the main text, for the sake of easier comparison of BNP-Track to other single particle tracking tools, we remove the time-dependence in emitter brightness and background flux to decrease the number of inferred quantities BNP-Track needs to estimate (where background flux, for instance, would need to be provided to competing tools). This can easily be done by modifying the equation above to

$$u_n^p = \tau_n^{\text{exps}} \left( CA^p + h \sum_{m=1}^{M} b^m Q_g^p(X_n^m, Y_n^m, Z_n^m) \right).$$

Here, $h$ and $C$ are just constant values for emitter brightness and background photon flux, respectively.

##### G.3 Emitter motion

To represent emitter motion, we consider dynamics consistent with free Brownian motion [43, 62, 51, 56, 31, 30, 7]. Accordingly, successive positions $X_n^m, Y_n^m, Z_n^m$ and $X_{n+1}^m, Y_{n+1}^m, Z_{n+1}^m$ or $X_{n-1}^m, Y_{n-1}^m, Z_{n-1}^m$ and $X_n^m, Y_n^m, Z_n^m$ along an emitter's trajectory are linked via independent Normal increments of zero mean and variance equal to $2D(T_{n+1} - T_n)$ or $2D(T_n - T_{n-1})$, respectively. In our model, $D$ indicated the emitters' diffusion coefficient.

For successive positions before a given time level, this leads to

$$\begin{aligned} X_n^m | X_{n+1}^m &\sim \text{Normal}\left(X_{n+1}^m, 2D(T_{n+1} - T_n)\right), \\ Y_n^m | Y_{n+1}^m &\sim \text{Normal}\left(Y_{n+1}^m, 2D(T_{n+1} - T_n)\right), \qquad n = 1, \dots, K^m - 1 \\ Z_n^m | Z_{n+1}^m &\sim \text{Normal}\left(Z_{n+1}^m, 2D(T_{n+1} - T_n)\right), \end{aligned}$$

while, for successive positions after a given time level, leads to

$$\begin{aligned} X_n^m | X_{n-1}^m &\sim \text{Normal}\left(X_{n-1}^m, 2D(T_n - T_{n-1})\right), \\ Y_n^m | Y_{n-1}^m &\sim \text{Normal}\left(Y_{n-1}^m, 2D(T_n - T_{n-1})\right), \qquad n = K^m + 1, \dots, N \\ Z_n^m | Z_{n-1}^m &\sim \text{Normal}\left(Z_{n-1}^m, 2D(T_n - T_{n-1})\right). \end{aligned}$$

Similar to all models' representing Brownian motion, our formulation leaves one position per trajectory unspecified. Namely, in our notation, the unspecified position of the $m^{\text{th}}$ emitter's trajectory corresponds to time level $n = K^m$ and needs to be modeled separately.

To model experiments under conditions that resemble stationary state, in which the statistics of the dynamical variables remain stable over the imaging course, we obtain the position left unspecified by

$$\begin{aligned} X_n^m &\sim \text{Uniform}_{[X_{\min}, X_{\max}]}, \\ Y_n^m &\sim \text{Uniform}_{[Y_{\min}, Y_{\max}]}, \qquad n = K^m \\ Z_n^m &\sim \text{Uniform}_{[Z_{\min}, Z_{\max}]} \end{aligned}$$

and the corresponding time level by

$$K^m \sim \text{Uniform}_{1:N}.$$

Overall, our representation of an emitter's motion is equivalent to the following Brownian probability density

$$\begin{aligned} p\left(X_{1:N}^m, Y_{1:N}^m, Z_{1:N}^m, K^m\right) = {} & \text{Uniform}_{[X_{\min}, X_{\max}]}\left(X_{K^m}^m\right) \prod_{n=1}^{N-1} \text{Normal}\left(X_{n+1} - X_n; 0, 2D(T_{n+1} - T_n)\right) \cdots \\ & \times \text{Uniform}_{[Y_{\min}, Y_{\max}]}\left(Y_{K^m}^m\right) \prod_{n=1}^{N-1} \text{Normal}\left(Y_{n+1} - Y_n; 0, 2D(T_{n+1} - T_n)\right) \cdots \\ & \times \text{Uniform}_{[Z_{\min}, Z_{\max}]}\left(Z_{K^m}^m\right) \prod_{n=1}^{N-1} \text{Normal}\left(Z_{n+1} - Z_n; 0, 2D(T_{n+1} - T_n)\right) \cdots \end{aligned}$$

$$\times \text{Uniform}_{1:N}(K^m)$$
$$\propto \prod_{n=1}^{N-1} \text{Normal}\left(X_{n+1} - X_n; 0, 2D(T_{n+1} - T_n)\right) \cdots$$
$$\times \prod_{n=1}^{N-1} \text{Normal}\left(Y_{n+1} - Y_n; 0, 2D(T_{n+1} - T_n)\right) \cdots$$
$$\times \prod_{n=1}^{N-1} \text{Normal}\left(Z_{n+1} - Z_n; 0, 2D(T_{n+1} - T_n)\right).$$

The proportionality above shows that our representation of the emitters' motion is consistent with studies characterizing single particle motion and diffusion coefficients by means of mean square displacements [7, 45, 51].

##### G.4 Extend BNP-Track to estimate multiple positions of one emitter within one frame

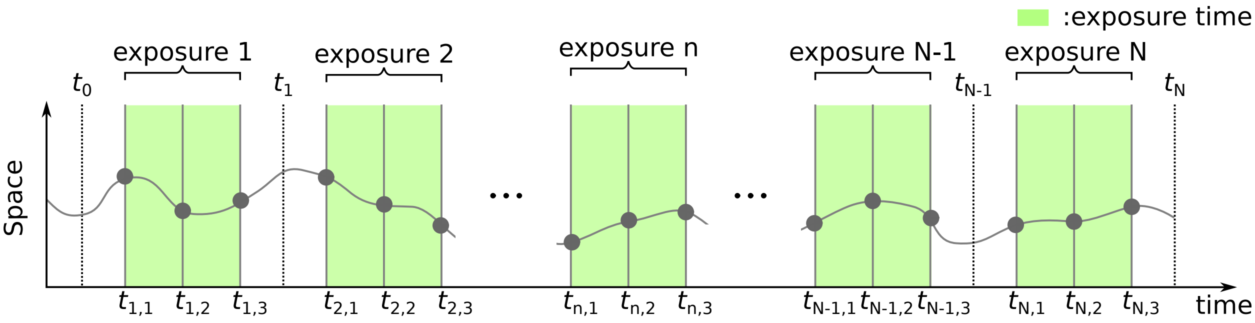

***Figure G.1:*** *How BNP-Track can be extended to consider intra-exposure emitter motion. In this figure, we show an example of estimating three positions for each emitter within each exposure.*

As a general tracking framework, BNP-Track, can be extended to estimate multiple positions of one emitter within one frame, see fig. G.1. This is important when emitters move appreciably as compared to a pixel dimension within one exposure resulting motion blur also termed aliasing, which distorts the PSF [21]. To address this issue, we can better approximate the first equation in appendix G using the trapezoidal rule. To be more specific, we can replace our simple quadrature rule

$$\int_{t_n^{\min}}^{t_n^{\max}} dt\, h(t) Q_g^p(X^m(t), Y^m(t), Z^m(t)) \approx \tau_n^{\text{exps}}\, h_n Q_g^p(X_n^m, Y_n^m, Z_n^m)$$

with a composite one as follows

$$\int_{t_n^{\min}}^{t_n^{\max}} dt\, h(t) Q_g^p(X^m(t), Y^m(t), Z^m(t))$$
$$\approx \frac{\tau_n^{\text{exps}}}{K-1} \sum_{k=1}^{K-1} \left[h_{n,k} Q_g^p(X_{n,k}^m, Y_{n,k}^m, Z_{n,k}^m) + h_{n,k+1} Q_g^p(X_{n,k+1}^m, Y_{n,k+1}^m, Z_{n,k+1}^m)\right].$$

Here, $K$ is the number of discretized time points in an exposure, and we define $h_{n,k} = h(t_{n,k})$, and $X_{n,k}^m = X^m(t_{n,k})$.

### H Model likelihood

In the preceding sections, we outlined our spatial and temporal frames of reference (appendices D and E), within which we define the various quantities of interest, and our mathematical formulation (appendices F and G), which establishes the relationships between those quantities. However, in an imaging experiment, most of the quantities of interest have unknown values. For example, in a typical superresolution experiment, the unknowns include

$$\theta = \left\{C_{1:N}, h_{1:N}, D, b^{1:M}, X^{1:M}_{1:N}, Y^{1:M}_{1:N}, Z^{1:M}_{1:N}, K^{1:M}\right\}.$$

As we explain in appendix C, our task is to infer these values through a posterior distribution that uses the images acquired.

In quantitative microscopy image measurements are provided in the form of image values $w^{1:P}_{1:N}$ obtained from the camera's array of pixel photosensors, which we label with $p = 1, \ldots, P$, and a sequence of exposures, which we label with $n = 1, \ldots, N$. Following the conventions introduced in appendices D, E and F.2, an image value $w^p_n$ probes the average photons $u^p_n$ incident on the region of space monitored by the $p^{\text{th}}$ pixel during the $n^{\text{th}}$ exposure. These correspond to

$$x^p_{\min} < x < x^p_{\max}, \qquad y^p_{\min} < y < y^p_{\max}, \qquad t^{\min}_n < t < t^{\max}_n.$$

Because of shot-noise and noisy camera read-out, the relation between $w^p_n$ and $u^p_n$ is stochastic and varies among different types of cameras. For scientific-grade cameras of the EMCCD type, which are commonly used in superresolution imaging [34, 28], the relationship is given by

$$w^p_n | u^p_n \sim \text{Normal}\left(\mu + \xi u^p_n, \upsilon + f\xi^2 u^p_n\right)$$

where $\mu$, $\upsilon$, $\xi$, and $f$ are the EMCCD's read-out offset, variance, overall gain, and excess noise factor [28], respectively. These values are reported in units of ADU, ADU$^2$, ADU/photon and $1$/photon, respectively. Because $\mu, \upsilon, \xi$, and $f$ are camera characteristics and not specimen characteristics, we obtain these values separately following the calibration protocol of appendix L.

Since $\mu, \upsilon, \xi$, and $f$ are specified via calibration before image processing, we use adjusted values

$$\bar{w}^p_n = \frac{w^p_n - \mu}{\xi}, \qquad \bar{\upsilon} = \frac{\upsilon}{\xi^2}$$

and cast our model's likelihood in its equivalent form

$$\bar{w}^p_n | u^p_n \sim \text{Normal}\left(u^p_n, \bar{\upsilon} + f u^p_n\right)$$

which allows for faster evaluation. With this convention, according to the definitions of table M.1, our likelihood's probability density takes the form

$$p(\bar{w}^p_n | u^p_n) = \frac{1}{\sqrt{2\pi(\bar{\upsilon} + f u^p_n)}} \exp\left(-\frac{1}{2}\frac{(\bar{w}^p_n - u^p_n)^2}{\bar{\upsilon} + f u^p_n}\right).$$

Finally, we may invoke a Laplace approximation [6, 27], according to

$$p(\bar{w}^p_n | u^p_n) \approx \frac{1}{\sqrt{2\pi(\bar{\upsilon} + f \bar{w}^p_n)}} \exp\left(-\frac{1}{2}\frac{(\bar{w}^p_n - u^p_n)^2}{\bar{\upsilon} + f u^p_n}\right) \propto \exp\left(-\frac{1}{2}\frac{(\bar{w}^p_n - u^p_n)^2}{\bar{\upsilon} + f u^p_n}\right)$$

to speed up the evaluation of our likelihood even further. Under the typical values encountered in a superresolution setting, such an approximation has negligible effect on our results.

### I Model priors

#### I.1 Nonparametric prior distributions

Our method is nonparametric and relies on indicator parameters denoted by $b^{1:M}$, which play a crucial role in our analysis. As we describe in METHODS, these parameters are model variables whose values are inferred concurrently with the other unknowns during processing.

To infer the values of $b^{1:M}$, we use independent Bernoulli priors with emitter-specific hyperparameters $q^{1:M}$, given by:

$$b^m | q^m \sim \text{Bernoulli}\left(q^m\right),$$

To avoid overfitting as the number of model emitters grows, we employ Beta hyper-priors of the form:

$$q^m \sim \text{Beta}\left(\frac{\gamma}{M}, 1 - \frac{\gamma}{M}\right)$$

on the Bernoulli weights $q^{1:M}$. With these priors and hyper-priors, our model remains well-defined at the limit $M \to \infty$, where it converges to a Beta-Bernoulli process [3, 12, 39, 42, 41]. The choice of $M$ has no interpretational side effects and only affects computation speed. Provided $M$ is sufficiently large, the precise value of $M$ has an insignificant influence on the results.

Because the prior and hyper-prior on each $b^m$ is independent of the others, we can combine them into a single distribution which takes the form

$$b^m \sim \text{Bernoulli}\left(\frac{\gamma}{M}\right)$$

and, because the total number of imaged emitters $B = \sum_{m=1}^{M} b^m$ is a derived variable, we can derive its prior statistics. In particular, as a sum of independent Bernoulli variables, the induced prior is

$$B \sim \text{Binomial}\left(M, \frac{\gamma}{M}\right)$$

which, at the limit $M \to \infty$, converges to a Poisson($\gamma$) random variable. Accordingly, $\gamma$ can be interpreted as the prior mean number of light emitters anticipated to contribute photons in the supplied images.

#### I.2 Parametric prior distributions

We have already described our choices for the nonparametric prior on the indicators $b^{1:M}$ as well as appropriate dynamical representations for the emitter positions $X_{1:N}^{1:M}$, $Y_{1:N}^{1:M}$, $Z_{1:N}^{1:M}$, and $K^{1:M}$. To complete our model, for the remaining unknowns $C_{1:N}$, $h_{1:N}$, and $D$ we make the following choices.

On the background photon fluxes $C_{1:N}$ and emitter photon emission rates $h_{1:N}$ we place independent Gamma priors

$$C_n \sim \text{Gamma}\left(A_n^C, \frac{C_n^{\text{ref}}}{A_n^C}\right),$$

$$h_n \sim \text{Gamma}\left(A_n^h, \frac{h_n^{\text{ref}}}{A_n^h}\right),$$

These depend on reference values $C_{1:N}^{\text{ref}}$ and $h_{1:N}^{\text{ref}}$ that set prior scales and provide units, as well as, on dimensionless parameters $A_{1:N}^C$ and $A_{1:N}^h$ that can be used to model prior confidence on the assigned reference values $C_{1:N}^{\text{ref}}$ and

$h^{\text{ref}}_{1:N}$ or lack thereof. For example, setting $A^C_n \gg 1$ restricts the background fluxes $C_{1:N}$ to attain values near $C^{\text{ref}}_n$ and similarly for $h_{1:N}$.

Finally, on the diffusion coefficient $D$ we place an Inverse-Gamma prior

$$D \sim \mathsf{InvGamma}\left(\alpha_D, (\alpha_D - 1)D_{\text{ref}}\right)$$

with, similarly, prior scale and units set by $D_{\text{ref}}$ and prior confidence set by $\alpha_D$.

#### J Summary of model equations

Our entire method for Bayesian nonparametric processing of imaging data is summarized in the following set of equations.

$$C_n \sim \text{Gamma}\left(A_n^C, C_n^{\text{ref}}/A_n^C\right), \qquad n = 1, \dots, N$$
$$h_n \sim \text{Gamma}\left(A_n^h, h_n^{\text{ref}}/A_n^h\right), \qquad n = 1, \dots, N$$

$$D \sim \text{InvGamma}\left(\alpha_D, (\alpha_D - 1)D_{\text{ref}}\right),$$

$$b^m \sim \text{Bernoulli}\left(\gamma/M\right), \qquad m = 1, \dots, M$$

$$K^m \sim \text{Uniform}_{1:N}, \qquad m = 1, \dots, M$$

$$X_n^m | K^m \sim \text{Uniform}_{[X_{\min}, X_{\max}]}, \qquad n = K^m \qquad m = 1, \dots, M$$
$$X_n^m | X_{n+1}^m, K^m, D \sim \text{Normal}\left(X_{n+1}^m, 2D(T_{n+1} - T_n)\right), \qquad n = 1, \dots, K^m - 1 \qquad m = 1, \dots, M$$
$$X_n^m | X_{n-1}^m, K^m, D \sim \text{Normal}\left(X_{n-1}^m, 2D(T_n - T_{n-1})\right), \qquad n = K^m + 1, \dots, N \qquad m = 1, \dots, M$$

$$Y_n^m | K^m \sim \text{Uniform}_{[Y_{\min}, Y_{\max}]}, \qquad n = K^m \qquad m = 1, \dots, M$$
$$Y_n^m | Y_{n+1}^m, K^m, D \sim \text{Normal}\left(Y_{n+1}^m, 2D(T_{n+1} - T_n)\right), \qquad n = 1, \dots, K^m - 1 \qquad m = 1, \dots, M$$
$$Y_n^m | Y_{n-1}^m, K^m, D \sim \text{Normal}\left(Y_{n-1}^m, 2D(T_n - T_{n-1})\right), \qquad n = K^m + 1, \dots, N \qquad m = 1, \dots, M$$

$$Z_n^m | K^m \sim \text{Uniform}_{[Z_{\min}, Z_{\max}]}, \qquad n = K^m \qquad m = 1, \dots, M$$
$$Z_n^m | Z_{n+1}^m, K^m, D \sim \text{Normal}\left(Z_{n+1}^m, 2D(T_{n+1} - T_n)\right), \qquad n = 1, \dots, K^m - 1 \qquad m = 1, \dots, M$$
$$Z_n^m | Z_{n-1}^m, K^m, D \sim \text{Normal}\left(Z_{n-1}^m, 2D(T_n - T_{n-1})\right), \qquad n = K^m + 1, \dots, N \qquad m = 1, \dots, M$$

$$w_n^p | C_n, h_n, b^{1:M}, X_n^{1:M}, Y_n^{1:M}, Z_n^{1:M} \sim \text{Normal}\left(\mu + \xi u_n^p, \upsilon + f\xi^2 u_n^p\right), \qquad n = 1, \dots, N, \qquad p = 1, \dots, P$$

The last equation may be substituted with its equivalent form,

$$\bar{w}_n^p | C_n, h_n, b^{1:M}, X_n^{1:M}, Y_n^{1:M}, Z_n^{1:M} \sim \text{Normal}\left(u_n^p, \bar{\upsilon} + f u_n^p\right), \qquad n = 1, \dots, N, \qquad p = 1, \dots, P$$

or approximated as described in appendix H.

In all equations, the parameters $C_n$, $h_n$, $b^{1:M}$, $X_n^{1:M}$, $Y_n^{1:M}$, and $Z_n^{1:M}$ are related to the image measurements by

$$u_n^p = \tau_n^{\text{exps}} \left( C_n A^p + h_n \sum_{m=1}^{M} b^m Q_g^p(X_n^m, Y_n^m, Z_n^m) \right).$$

### K Evaluation and interpretation of the model posterior

Our image analysis method, BNP-Track, is fully contained in the statistical model whose equations are listed in appendix J. We have adapted these equations to meet the particular imaging conditions of superresolution microscopy as well as to facilitate the estimation of the variables of typical interest in a biological, biochemical, or biophysical experiment [34] that we detailed in appendices D to G.

As we explain in appendix C, we encode our model's equations and an experiment's measured images in a posterior probability distribution [9, 53, 59, 18]. This distribution assimilates into our analysis the information supplied by the form of the model itself (e.g. information such as parameter ranges or relations among the parameters) and the information supplied by the experimental data, which takes the form of image values reported by the camera devices used.

Our posterior distribution, in its complete form, is

$$p\Big(\underbrace{C_{1:N}, h_{1:N}, D, b^{1:M}, X_{1:N}^{1:M}, Y_{1:N}^{1:M}, Z_{1:N}^{1:M}, K^{1:M}}_{\text{unknowns}} \Big| \underbrace{w_{1:N}^{1:P}}_{\text{data}}\Big).$$

For convenience, we define a single variable $\theta$ that gathers all unknowns, including $C_{1:N}$, $h_{1:N}$, $D$, $b^{1:M}$, $X_{1:N}^{1:M}$, $Y_{1:N}^{1:M}$, $Z_{1:N}^{1:M}$, and $K^{1:M}$. We also use $W$ to represent the image measurement, which is denoted as $w_{1:N}^{1:P}$. With this notation, the posterior distribution can be compactly expressed as $p(\theta|W)$.

The posterior $p(\theta|W)$ quantifies, in an absolute scale, the consistency between any given assignment of specific values to the unknowns, e.g. an assignment such as $C_1 = 5 \times 10^4$ photons/s/$\mu$m$^2$, $h_1 = 6 \times 10^4$ photons/s, $D = 0.11$ $\mu$m$^2$/s and so on, with the measured images. In principle, with $p(\theta|W)$ we can precisely quantify how probable is for the assigned values to have generated the recorded images; high posterior corresponds to highly probable value assignments and, vice versa, low posterior corresponds to highly improbable assignments. Nevertheless, due to the nonparametric prior and the non-trivial relations among our unknowns, an exhaustive quantification of all possible value assignments via greedy computations is impossible. This is because, at the limit $M \to \infty$, our infinite number of unknowns $b^{1:M}$, $X_{1:N}^{1:M}$, $Y_{1:N}^{1:M}$, $Z_{1:N}^{1:M}$, $K^{1:M}$, and the non-linear dependencies underlying $u_{1:N}^{1:P}$ lead to intractable formulas.

Instead, we use the posterior $p(\theta|W)$ in a different, but statistically equivalent, manner. In particular, instead of seeking to quantify exhaustively every possible parameter value assignment, the vast majority of which is highly inconsistent with the measured images, we compute only those value assignments that are most consistent with the images [18]. This way we quantify value assignment in a relative scale. Practically, as we illustrate in appendix C, we generate computationally random assignments in such a way that most probable values occur with higher frequency than the others. As we demonstrate in RESULTS and fig. C.1, we can derive any statistic of interest by those assignments which, to avoid confusion, from now on we will call posterior samples.

Next, we describe the steps necessary to generate and handle posterior samples.

#### K.1 Evaluation

For clarity, we denote the posterior samples we seek to generate with superscripts $(i)$. Specifically, these are $\theta^{(0)}, \theta^{(1)}, \theta^{(2)}, \dots, \theta^{(i)}, \theta^{(i+1)}, \dots$. To generate these samples, we develop a Markov chain Monte Carlo scheme [47, 35]. Namely, we use pseudo-random computer simulations to advance from one sample to the next one. For this, we apply a custom Gibbs sampling scheme [18, 47, 48, 35].

Specifically, we initialize values in $\theta^{(0)}$ for all variables by sampling from their respective priors. Subsequently, to advance from $\theta^{(i)}$ to $\theta^{(i+1)}$, our scheme proceeds in steps during which we update only a designated block of parameters. In detail, we update the parameters in blocks by interlacing, in random order, the following six steps:

1. We update the variables $C_{1:N}$. This step, involves sampling from the conditional distribution
$$p\Big(C_{1:N}\Big|h_{1:N}, D, b^{1:M}, X_{1:N}^{1:M}, Y_{1:N}^{1:M}, Z_{1:N}^{1:M}, K^{1:M}, w_{1:N}^{1:P}\Big).$$
2. We update the variables $C_{1:N}$ and $h_{1:N}$. This step, involves sampling from the conditional distribution
$$p\Big(C_{1:N}, h_{1:N}\Big|D, b^{1:M}, X_{1:N}^{1:M}, Y_{1:N}^{1:M}, Z_{1:N}^{1:M}, K^{1:M}, w_{1:N}^{1:P}\Big).$$
3. We update the variables $b^{1:M}, X_{1:N}^{1:M}, Y_{1:N}^{1:M}, Z_{1:N}^{1:M}, K^{1:M}$. This step, involves sampling from the conditional distribution
$$p\Big(b^{1:M}, X_{1:N}^{1:M}, Y_{1:N}^{1:M}, Z_{1:N}^{1:M}, K^{1:M}\Big|C_{1:N}, h_{1:N}, D, w_{1:N}^{1:P}\Big).$$
4. We update the variables $C_{1:N}$ and $b^{1:M}, X_{1:N}^{1:M}, Y_{1:N}^{1:M}, Z_{1:N}^{1:M}, K^{1:M}$. This step, involves sampling from the conditional distribution
$$p\Big(C_{1:N}, b^{1:M}, X_{1:N}^{1:M}, Y_{1:N}^{1:M}, Z_{1:N}^{1:M}, K^{1:M}\Big|h_{1:N}, D, w_{1:N}^{1:P}\Big).$$
5. We update the variables $C_{1:N}$ and $X_{1:N}^{1:M}, Y_{1:N}^{1:M}, Z_{1:N}^{1:M}, K^{1:M}$. This step, involves sampling from the conditional distribution
$$p\Big(C_{1:N}, X_{1:N}^{1:M}, Y_{1:N}^{1:M}, Z_{1:N}^{1:M}, K^{1:M}\Big|h_{1:N}, b^{1:M}, D, w_{1:N}^{1:P}\Big).$$
6. We update the variables $X_{1:N}^{1:M}, Y_{1:N}^{1:M}, Z_{1:N}^{1:M}, K^{1:M}$. This step, involves sampling from the conditional distribution
$$p\Big(X_{1:N}^{1:M}, Y_{1:N}^{1:M}, Z_{1:N}^{1:M}, K^{1:M}\Big|C_{1:N}, h_{1:N}, b^{1:M}, D, w_{1:N}^{1:P}\Big).$$

For all steps we use customized Metropolis-Hastings samplers [48, 47, 18, 35]. Our samplers are purpose-built and use techniques based on slice-sampling [37, 38, 36], or multiplicative random walks [15]. We evaluate our posterior with a fixed and finite total number of model emitters $M \gg 1$. As we explain in appendix I.1, this introduces an approximation in our analysis with an insignificant error [40, 3].

Although our computational scheme is mathematically valid, its performance can be improved considerably if it is combined with simulated annealing [26, 57, 8, 23]. Annealing can speed up the convergence rate, reducing processing time which, for image data of typical sizes, may be slow. In our sampler, we introduce an annealing factor $F^{(i)}$ and implement simulated annealing by replacing the EMCCD's excess noise factor $f$ in our likelihood by the product $F^{(i)}f$. The annealing factor $F^{(i)} \geq 1$ is set to a large value in the beginning of our Gibbs iterations and gradually reduces to $1$, for instance such as in

$$F^{(i)} = 1 + (F^{(0)} - 1)\max\left(0, 1 - i/i_{\text{ref}}\right)^2$$

where $F^{(0)}$ indicates the initial value, $i_{\text{ref}}$ the reduction speed, and $i$ is the iteration number.

##### K.2 Interpretation

Once a sequence of posterior samples $\theta^{(1)}, \theta^{(2)}, \dots$ is obtained by the successive repetition of the Gibbs sampling scheme, we may compute any statistic directly related to the sampled variables [59, 18, 48, 47, 35].

As with every Markov chain Monte Carlo method [59, 18, 48, 47, 35], the accuracy of the computed statistics is drastically improved if, from the entire chain of posterior samples, we first discard an initial burn-in portion. To identify this portion we use batching [48]. Specifically, once simulated annealing terminates at iteration $i = i_{\text{ref}}$, we divide the remaining sequence into three successive batches each covering 1/3 of successive samples in the chain. For each of the last two batches we compute the statistics of interest and compare. If these agree, we use either of them as our estimate; however, if they do not agree, we expand our initial chain by computing additional posterior samples and repeat until the statistics of the terminal two batches match.

Similarly, we may also compute statistics of interest also of derived quantities that depend upon the sampled variables [48, 35]. One such quantity is the total number $B$ of light emitters that contribute photons to the measured images. Since $B$ depends upon the sampled indicators $b^{1:M}$, its posterior samples are readily obtained by the sum $B = \sum_{m=1}^{M} b^m$.

Unlike $B$, however, which is uniquely determined by the sampled variables, statistics that are sensitive to the labeling of the emitters, i.e. statistics that require distinctive labels $m$ assigned to each model emitter, may not be determined uniquely by the sampled variables. This is a common characteristic shared by Bayesian nonparametric methods [44, 49, 54] and reflects the fact that, both a priori and a posteriori, all emitters are equivalent in the sense that there is no preference for a particular labeling out of all $M!$ possible ones. For this reason, following our Markov chain Monte Carlo computations, we relabel the emitters in our posterior samples such that they maintain fixed labels. Essentially, from the $M!$ equivalent posterior modes that our posterior allows for, with our relabeling approach, we chose methodologically only one to base our estimates upon. We explain our relabeling strategy below.

In any posterior sample $\theta$, the variables requiring relabeling are $b^{1:M}, X_{1:N}^{1:M}, Y_{1:N}^{1:M}, Z_{1:N}^{1:M}, K^{1:M}$. For clarity, we denote their relabeled counterparts with $\dot{\theta}$ and $\dot{b}^{1:M}, \dot{X}_{1:N}^{1:M}, \dot{Y}_{1:N}^{1:M}, \dot{Z}_{1:N}^{1:M}, \dot{K}^{1:M}$, respectively. To obtain these, we first choose a pivot sample which we select out of the computed ones $\theta^{(i)}$ that remain after burn-in removal. Our pivot is the posterior sample that corresponds to the highest posterior $p(\theta|W)$. For clarity, we denote the pivot with $\tilde{\theta}$ and its variables with $\tilde{b}^{1:M}, \tilde{X}_{1:N}^{1:M}, \tilde{Y}_{1:N}^{1:M}, \tilde{Z}_{1:N}^{1:M}, \tilde{K}^{1:M}$. Subsequently, for each available $\theta^{(i)}$ we form all $M!$ possible samples $\ddot{\theta}_k^{(i)}$ through the permutations of the labels and for $\dot{\theta}^{(i)}$ we select the sample $\ddot{\theta}_k^{(i)}$ that is most similar to the pivot.

Our comparison with the pivot is based on the similarity metric

$$\mathcal{D}\left(\ddot{\theta}; \tilde{\theta}\right) = \sum_{m=1}^{M} \mathcal{R}_{\ddot{g}}\left(\ddot{b}^m, \ddot{X}_{1:N}^m, \ddot{Y}_{1:N}^m, \ddot{Z}_{1:N}^m; \tilde{b}^m, \tilde{X}_{1:N}^m, \tilde{Y}_{1:N}^m, \tilde{Z}_{1:N}^m\right)$$

which, in turn, depends additively on the metric

$$\mathcal{R}_{\ddot{g}}\left(\ddot{b}, \ddot{X}_{1:N}, \ddot{Y}_{1:N}, \ddot{Z}_{1:N}; \tilde{b}, \tilde{X}_{1:N}, \tilde{Y}_{1:N}, \tilde{Z}_{1:N}\right) = \sum_{p=1}^{P} \sum_{n=1}^{N} \tau_n^{\text{exps}} h_n \left| \ddot{b}\, Q_{\ddot{g}}^p(\ddot{X}_n, \ddot{Y}_n, \ddot{Z}_n) - \tilde{b}\, Q_{\ddot{g}}^p(\tilde{X}_n, \tilde{Y}_n, \tilde{Z}_n) \right|,$$

that compares $\ddot{\theta}$ and $\tilde{\theta}$ emitterwise based on the respective images. From these two, the former metric compares the entire population of emitters; while, the latter metric compares individual emitters. Because the population metric $\mathcal{D}\left(\ddot{\theta}; \tilde{\theta}\right)$ depends additively on the emitter metric $\mathcal{R}_{\ddot{g}}\left(\ddot{b}, \ddot{X}_{1:N}, \ddot{Y}_{1:N}, \ddot{Z}_{1:N}; \tilde{b}, \tilde{X}_{1:N}, \tilde{Y}_{1:N}, \tilde{Z}_{1:N}\right)$, selecting the optimum $\dot{\theta}^{(i)}$ out of $\ddot{\theta}_{1:M!}^{(i)}$ reduces to a linear assignment problem that can be solved efficiently [14, 11], for instance through the Hungarian algorithm, without explicitly forming the permutations $\ddot{\theta}_{1:M!}^{(i)}$, which is impractical.

### L Calibration

#### L.1 Camera read-out

For cameras of the EMCCD type, agreement with the signal-to-noise specifications [29, 28, 24] requires a fixed excess noise factor $f = 2/\text{photon}$. However, the other camera parameters, e.g. $\mu$, $\upsilon$ and $\xi$, are device and configuration dependent [24]. For this reason, their values must be calibrated separately for each camera and imaging configuration adopted. Below we describe a standard calibration procedure.

The values of readout offset $\mu$ and variance $\upsilon$ can be evaluated with dark images, i.e. images under no photon flux such as those obtained with the camera's shutter closed [29, 24]. We denote with $\omega_{1:N}^{0,1:P}$ such recorded images. Specifically, for these $u_n^{0,p} = 0$, and so our model reduces to

$$\omega_n^{0,p} \sim \text{Normal}\left(\mu, \upsilon\right).$$

For a sufficiently large number of exposures $N$, offset $\mu$ and variance $\upsilon$ can be recovered by the sample mean and variance, respectively

$$\mu = \frac{1}{NP}\sum_{n=1}^{N}\sum_{p=1}^{P}\omega_n^{0,p}, \qquad \upsilon = \frac{1}{NP-1}\sum_{n=1}^{N}\sum_{p=1}^{P}\left(\omega_n^{0,p} - \mu\right)^2.$$

The value of the gain $\xi$ can be evaluated based on static images, i.e. images under constant photon flux such as those obtained when imaging an illuminated empty sample in plain buffer or other optically homogenous media. We denote with $\omega_{1:N}^{k,1:P}$ such recorded images, and use super-scripts $k = 1, \ldots, K$ to denote different illumination levels. Specifically, for such images $u_n^{k,p} = C_{\text{static}}^{p,k} A^p \tau_n^{\text{exps}}$ where $C_{\text{static}}^{p,k}$ is the photon flux achieved at the $k^{\text{th}}$ illumination level, and so our model reduces to

$$\omega_n^{k,p} \sim \text{Normal}\left(\mu + \xi u_n^{k,p}, \upsilon + f\xi^2 u_n^{k,p}\right)$$

Accordingly, for each illumination level, each pixel's recordings mean and variance across exposures are given by $\mu^{k,p} = u_n^{k,p}\xi + \mu$ and $\upsilon^{k,p} = u_n^{k,p} f\xi^2 + \upsilon$, respectively. Consequently, they are related to each other by $\upsilon^{k,p} - \upsilon = f\xi\left(\mu^{k,p} - \mu\right)$. This relation can be used to obtain the value of $\xi$ through least squares estimation [18, 25]. The result is

$$\xi = \frac{1}{f}\frac{\sum_{k=1}^{K}\sum_{n=1}^{N}\left(\upsilon^{k,p} - \upsilon\right)\left(\mu^{k,p} - \mu\right)}{\sum_{k=1}^{K}\sum_{n=1}^{N}\left(\mu^{k,p} - \mu\right)^2},$$

where the mean and variance of each pixel's recordings are recovered by the sample mean and variance, respectively

$$\mu^{k,p} = \frac{1}{N}\sum_{n=1}^{N}\omega_n^{k,p}, \qquad \upsilon^{k,p} = \frac{1}{N-1}\sum_{n=1}^{N}\left(\omega_n^{k,p} - \mu\right)^2.$$

#### L.2 Point spread function and illumination profile

For the calibration of the point spread function, suspended fiducial markers [17] can be imaged at multiple stages as graphically illustrated in fig. L.1. For clarity, we denote with $\omega_{1:N}^{*,1:P}$ the images obtained with corresponding stage displacement $d_{1:N}^{\text{stg}}$ and, to proceed, we denote with $\bar{X}^m, \bar{Y}^m, \bar{Z}^m$ the emitters' positions in a laboratory's frame of reference. Further, we assume that the latter is oriented such that, when the stage is displaced by $d$, the emitters are imaged at $X^m = \bar{X}^m, Y^m = \bar{Y}^m, Z^m = \bar{Z}^m + d$ with respect to the object plane which is our common frame

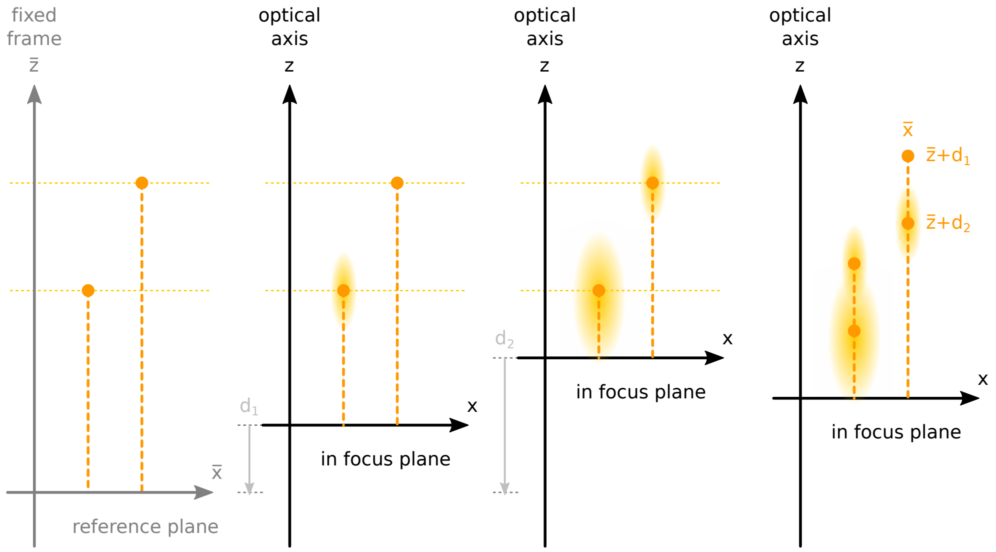

***Figure L.1:*** *Configuration of sample space and stage displacement used in the calibration of the point spread function. To increase clarity, here we depict only two of the three spacial coordinates. Suspended fiducial markers, immobile with respect to the laboratory's frame of reference (left panel), are imaged at different stages (middle panels). Image analysis allows for characterization and reconstruction of the point spread function along all spatial dimensions (right panel).*

of reference. Therefore, in the $n^{\text{th}}$ image, the emitters are located at $X_n^m = \bar{X}^m, Y_n^m = \bar{Y}^m, Z_n^m = \bar{Z}^m + d_n^{\text{stg}}$. We illustrate this convention in fig. L.1.

Assuming no emitter motion, the average number of incident photons in our calibration model reduces to

$$u_n^p = \tau_n^{\text{exps}} \left( C_n A^p + h \sum_{m=1}^{M} b^m \iint_{x_{\min}^p, y_{\min}^p}^{x_{\max}^p, y_{\max}^p} dx dy \; G_g(x, y; X_n^m, Y_n^m, Z_n^m) \right).$$

Under these assumptions, image analysis for the calibration of the point spread function can be performed with the model

$$\begin{aligned}
g_j &\sim \text{InvGamma}\left(\alpha_g, \alpha_g - 1\right), & j &= 1, 2 \\
C_n &\sim \text{Gamma}\left(\alpha_C, C_{\text{ref}}/\alpha_C\right), & n &= 1, \dots, N \\
h &\sim \text{Gamma}\left(\alpha_h, h_{\text{ref}}/\alpha_h\right) \\
b^m &\sim \text{Bernoulli}\left(\gamma/M\right), & m &= 1, \dots, M \\
\bar{X}^m &\sim \text{Uniform}_{[X_{\min}, X_{\max}]}, & m &= 1, \dots, M \\
\bar{Y}^m &\sim \text{Uniform}_{[Y_{\min}, Y_{\max}]}, & m &= 1, \dots, M \\
\bar{Z}^m &\sim \text{Uniform}_{[Z_{\min}, Z_{\max}]}, & m &= 1, \dots, M
\end{aligned}$$

$$\omega_n^{*,p} | g_{1:3}, C_n, h, b^{1:M}, \bar{X}^{1:M}, \bar{Y}^{1:M}, \bar{Z}^{1:M} \sim \text{Normal}\left(\mu + \xi u_n^p, \upsilon + f\xi^2 u_n^p\right), \quad n = 1, \dots, N, \quad p = 1, \dots, P$$

which yields estimates for $g_1$ and $g_2$. The estimation relies on the marginal posterior $p(g_1, g_2 | \omega_{1:N}^{*,1:P})$ which can be evaluated through Markov chain Monte Carlo sampling similar to the scheme we described in appendix K.

### M Definitions

***Table M.1:*** *Summary of probability distributions. In this table $\Gamma(\cdot)$ and $\mathrm{B}(\cdot,\cdot)$ denote the Gamma and Beta functions, respectively [1]. Further, $x!$ denotes the factorial of $x$ and $\delta_x(\cdot)$ denotes the Dirac delta centered at $x$.*

| Probability distribution | Variable | Values | Probability density |
|---|---|---|---|
| Normal$(\mu, v)$ | $x$ | from $-\infty$ to $+\infty$ | $\frac{1}{\sqrt{2\pi v}} \exp\left(-\frac{(\mu-x)^2}{2v}\right)$ |
| Gamma$(\phi, \beta)$ | $w$ | from $0$ to $\infty$ | $\frac{1}{\beta\Gamma(\phi)} \left(\frac{w}{\beta}\right)^{\phi-1} \exp\left(-\frac{w}{\beta}\right)$ |
| InvGamma$(\phi, \beta)$ | $d$ | from $0$ to $\infty$ | $\frac{1}{\beta\Gamma(\phi)} \left(\frac{\beta}{d}\right)^{\phi+1} \exp\left(-\frac{\beta}{d}\right)$ |
| Uniform$_{[x,y]}$ | $z$ | from $x$ to $y$ | $\frac{1}{y-x}$ |
| Beta$(\alpha, \beta)$ | $q$ | from $0$ to $1$ | $\frac{1}{\mathrm{B}(\alpha,\beta)} q^{\alpha-1}(1-q)^{\beta-1}$ |
| Bernoulli$(q)$ | $b$ | $0$ or $1$ | $q\delta_1(b) + (1-q)\delta_0(b)$ |
| Binomial$(m, r)$ | $b$ | $0$ or $1$ or $2$ or $\ldots$ | $\sum_{k=0}^{\infty} \frac{m!}{k!(m-k)!} r^k (1-r)^{m-k} \delta_k(b)$ |
| Poisson$(\gamma)$ | $b$ | $0$ or $1$ or $2$ or $\ldots$ | $\sum_{k=0}^{\infty} \frac{\gamma^k}{k!} e^{-\gamma} \delta_k(b)$ |
| Uniform$_{1:N}$ | $k$ | from $1$ to $N$ | $\frac{1}{N}$ |

#### References

[1] M. Abramowitz and I. A. Stegun. Handbook of mathematical functions with formulas, graphs, and mathematical tables, volume 55. US Government printing office, 1948.

[2] F. Aguet. Super-Resolution Fluorescence Microscopy Based on Physical Models. EPFL thesis no. 4418 (2009), 209 p., Swiss Federal Institute of Technology Lausanne (EPFL), 2009.

[3] L. Al Labadi and M. Zarepour. On approximations of the beta process in latent feature models: Point processes approach. Sankhya A, 80(1):59, 2018.

[4] K. Atkinson and W. Han. Theoretical numerical analysis, volume 39. Springer, 2005.

[5] D. W. Ball. Field guide to spectroscopy, volume 8. SPIE Press Bellingham, Washington, 2006.

[6] C. M. Bender, S. Orszag, and S. A. Orszag. Advanced mathematical methods for scientists and engineers I: Asymptotic methods and perturbation theory, volume 1. Springer Science & Business Media, 1999.

[7] H. C. Berg. Random walks in biology. In Random Walks in Biology. Princeton University Press, 2018.

[8] D. Bertsimas and J. Tsitsiklis. Simulated annealing. Statistical science, 8(1):10, 1993.

[9] C. M. Bishop. Pattern recognition and machine learning. Springer, 2006.

[10] M. Born and E. Wolf. Principles of optics: electromagnetic theory of propagation, interference and diffraction of light. Elsevier, 2013.

[11] R. Burkard, M. Dell'Amico, and S. Martello. Assignment Problems. Society for Industrial and Applied Mathematics, 2012.

[12] Y. Cheng, D. Li, and W. Jiang. The exact inference of beta process and beta bernoulli process from finite observations. Comput. Model. Eng. Sci., 121(1):49, 2019.

[13] H. Deschout, K. Neyts, and K. Braeckmans. The influence of movement on the localization precision of sub-resolution particles in fluorescence microscopy. J. Biophotonics, 5(1):97, 2012.

[14] I. S. Duff and J. Koster. On algorithms for permuting large entries to the diagonal of a sparse matrix. SIAM J. Matrix Anal. Appl., 22(4):973, 2001.

[15] S. Dutta. Multiplicative random walk metropolis-hastings on the real line. Sankhya B, 74(2):315, 2012.

[16] G. R. Fowles. Introduction to modern optics. Courier Corporation, 1989.

[17] C. Gell, M. Berndt, J. Enderlein, and S. Diez. Tirf microscopy evanescent field calibration using tilted fluorescent microtubules. J. Microsc., 234(1):38, 2009.

[18] A. Gelman, J. B. Carlin, H. S. Stern, D. B. Dunson, A. Vehtari, and D. B. Rubin. Bayesian data analysis. CRC press, 3rd edition, 2013.

[19] S. F. Gibson and F. Lanni. Experimental test of an analytical model of aberration in an oil-immersion objective lens used in three-dimensional light microscopy. JOSA A, 8(10):1601, 1991.

[20] J. W. Goodman. Introduction to Fourier optics. Roberts and Company Publishers, 2005.

[21] J. W. Goodman. Introduction to Fourier optics. Roberts and Company publishers, 2005.

[22] E. Hairer, C. Lubich, and G. Wanner. Geometric numerical integration: structure-preserving algorithms for ordinary differential equations, volume 31. Springer Science & Business Media, 2006.

[23] B. Hajek. Cooling schedules for optimal annealing. Math. Oper. Res., 13(2):311, 1988.

[24] K. B. Harpsøe, M. I. Andersen, and P. Kjægaard. Bayesian photon counting with electron-multiplying charge coupled devices (EMCCDs). Astron. Astrophys., 537:A50, 2012.

[25] T. Hastie, R. Tibshirani, J. H. Friedman, and J. H. Friedman. The elements of statistical learning: data mining, inference, and prediction, volume 2. Springer, 2009.

[26] D. Henderson, S. H. Jacobson, and A. W. Johnson. The theory and practice of simulated annealing. In Handbook of metaheuristics, page 287. Springer, 2003.

[27] E. J. Hinch. Perturbation Methods. Cambridge Texts in Applied Mathematics. Cambridge University Press, 1991.

[28] M. Hirsch, R. J. Wareham, M. L. Martin-Fernandez, M. P. Hobson, and D. J. Rolfe. A stochastic model for electron multiplication charge-coupled devices – from theory to practice. PLoS One, 8(1):1, 2013.

[29] F. Huang, T. M. P. Hartwich, F. E. Rivera-Molina, Y. Lin, W. C. Duim, J. J. Long, P. D. Uchil, J. R. Myers, M. A. Baird, W. Mothes, M. W. Davidson, D. Toomre, and J. Bewersdorf. Video-rate nanoscopy using scmos camera–specific single-molecule localization algorithms. Nat. Methods, 10(7):653, 2013.

[30] S. Jazani, I. Sgouralis, and S. Pressé. A method for single molecule tracking using a conventional single-focus confocal setup. J. Chem. Phys., 150(11):114108, 2019.

[31] S. Jazani, I. Sgouralis, O. M. Shafraz, M. Levitus, S. Sivasankar, and S. Pressé. An alternative framework for fluorescence correlation spectroscopy. Nat. Commun., 10(1):3662, 2019.

[32] Z. Kilic, I. Sgouralis, W. Heo, K. Ishii, T. Tahara, and S. Pressé. Extraction of rapid kinetics from smfret measurements using integrative detectors. Cell Rep. Phys. Sci., 2(5):100409, 2021.

[33] J. R. Lakowicz. Principles of fluorescence spectroscopy. Springer science & business media, 2013.

[34] A. Lee, K. Tsekouras, C. Calderon, C. Bustamante, and S. Pressé. Unraveling the thousand word picture: An introduction to super-resolution data analysis. Chem. Rev., 117(11):7276, 2017.

[35] J. S. Liu. Monte Carlo strategies in scientific computing. Springer Science & Business Media, 2008.

[36] I. Murray, R. Adams, and D. MacKay. Elliptical slice sampling. In Proceedings of the thirteenth international conference on artificial intelligence and statistics, page 541. JMLR Workshop and Conference Proceedings, 2010.

[37] R. M. Neal. Slice sampling. Ann. Stat., 31(3):705, 2003.

[38] R. Nishihara, I. Murray, and R. P. Adams. Parallel mcmc with generalized elliptical slice sampling. J. Mach. Learn. Res., 15(1):2087, 2014.

[39] J. Paisley, D. Blei, and M. Jordan. Stick-breaking beta processes and the poisson process. In Artificial Intelligence and Statistics, page 850. PMLR, 2012.

[40] J. Paisley and L. Carin. Nonparametric factor analysis with beta process priors. In Proceedings of the 26th annual international conference on machine learning, pages 777–784, 2009.

[41] J. Paisley, L. Carin, and D. Blei. Variational inference for stick-breaking beta process priors. In Proceedings of the 28th international conference on machine learning, 2011.

[42] J. W. Paisley, A. K. Zaas, C. W. Woods, G. S. Ginsburg, and L. Carin. A stick-breaking construction of the beta process. In ICML, 2010.

[43] G. A. Pavliotis. Stochastic processes and applications: diffusion processes, the Fokker-Planck and Langevin equations, volume 60. Springer, 2014.

[44] K. Puolamäki and S. Kaski. Bayesian solutions to the label switching problem. In International Symposium on Intelligent Data Analysis, page 381. Springer, 2009.

[45] H. Qian, M. P. Sheetz, and E. L. Elson. Single particle tracking. analysis of diffusion and flow in two-dimensional systems. Biophys. J., 60(4):910, 1991.

[46] A. Quarteroni, R. Sacco, and F. Saleri. Numerical mathematics, volume 37. Springer Science & Business Media, 2010.

[47] C. Robert and G. Casella. Monte Carlo statistical methods. Springer Science & Business Media, 2013.

[48] C. P. Robert, G. Casella, and G. Casella. Introducing monte carlo methods with R, volume 18. Springer, 2010.

[49] C. E. Rodríguez and S. G. Walker. Label switching in bayesian mixture models: Deterministic relabeling strategies. Journal of Computational and Graphical Statistics, 23(1):25, 2014.

[50] B. E. Saleh and M. C. Teich. Fundamentals of photonics. john Wiley & sons, 2019.

[51] T. Schlick. Molecular modeling and simulation: an interdisciplinary guide, volume 21. Springer Science & Business Media, 2010.

[52] A. E. Siegman. Lasers. University Science Books, 1986.

[53] D. Sivia and J. Skilling. Data analysis: a Bayesian tutorial. OUP Oxford, 2006.

[54] M. Stephens. Dealing with label switching in mixture models. J. R. Stat. Soc., B: Stat. Methodol., 62(4):795, 2000.

[55] L. Tao and C. Nicholson. The three-dimensional point spread functions of a microscope objective in image and object space. J. Microsc., 178(3):267, 1995.

[56] M. Tavakoli, S. Jazani, I. Sgouralis, O. M. Shafraz, S. Sivasankar, B. Donaphon, M. Levitus, and S. Pressé. Pitching single-focus confocal data analysis one photon at a time with bayesian nonparametrics. Phys. Rev. X, 10:011021, 2020.

[57] P. J. Van Laarhoven and E. H. Aarts. Simulated annealing. In Simulated annealing: Theory and applications, page 7. Springer, 1987.

[58] D. G. Voelz. Computational fourier optics: a MATLAB tutorial, volume 534. SPIE press Bellingham, Washington, 2011.

[59] U. von Toussaint. Bayesian inference in physics. Rev. Mod. Phys., 83:943, 2011.

[60] M. S. Wartak. Computational photonics: an introduction with MATLAB. Cambridge University Press, 2013.

[61] B. Zhang, J. Zerubia, and J.-C. Olivo-Marin. Gaussian approximations of fluorescence microscope point-spread function models. Appl. Opt., 46(10):1819, 2007.

[62] R. Zwanzig. Nonequilibrium statistical mechanics. Oxford University Press, 2001.
